# Cell differentiation controls iron assimilation in a choanoflagellate

**DOI:** 10.1101/2024.05.25.595918

**Authors:** Fredrick Leon, Jesus M. Espinoza-Esparza, Vicki Deng, Maxwell C. Coyle, Sarah Espinoza, David S. Booth

**Affiliations:** Chan Zuckerberg Biohub & Department of Biochemistry and Biophysics, University of California, San Francisco School of Medicine San Francisco, CA 94143; Howard Hughes Medical Institute & Department of Molecular and Cell Biology University of California, Berkeley Berkeley, CA 94720; Department of Molecular Biosciences University of Texas, Austin Austin, TX 78712; Department of Molecular and Cellular Biology Harvard University, Cambridge, MA 02138

## Abstract

Marine microeukaryotes have evolved diverse cellular features that link their life histories to surrounding environments. How those dynamic life histories intersect with the ecological functions of microeukaryotes remains a frontier to understand their roles in essential biogeochemical cycles^1,2^. Choanoflagellates, phagotrophs that cycle nutrients through filter feeding, provide models to explore this intersection, for many choanoflagellate species transition between life history stages by differentiating into distinct cell types^3–6^. Here we report that cell differentiation in the marine choanoflagellate *Salpingoeca rosetta* endows one of its cell types with the ability to utilize insoluble ferric colloids for improved growth through the expression of a cytochrome b561 iron reductase (*cytb561a*). This gene is an ortholog of the mammalian duodenal cytochrome b561 (*DCYTB*) that reduces ferric cations prior to their uptake in gut epithelia^7^ and is part of an iron utilization toolkit that choanoflagellates and their closest living relatives, the animals, inherited from a last common eukaryotic ancestor. In a database of oceanic metagenomes^8,9^, the abundance of *cytb561a* transcripts from choanoflagellates positively correlates with upwellings, which are a major source of ferric colloids in marine environments^10^. As this predominant form of iron^11,12^ is largely inaccessible to cell-walled microbes^13,14^, choanoflagellates and other phagotrophic eukaryotes may serve critical ecological roles by first acquiring ferric colloids through phagocytosis and then cycling this essential nutrient through iron utilization pathways^13–15^. These findings provide insight into the ecological roles choanoflagellates perform and inform reconstructions of early animal evolution where functionally distinct cell types became an integrated whole at the origin of animal multicellularity^16–22^.

## Text

### Thecates display a different transcriptome profile than other cell types

Choanoflagellates transition between different stages of their life history by differentiating into distinct cell types^3–5^. In the emerging model choanoflagellate *S. rosetta*, diverse environmental cues promote the transitions between different types of cells^6^. Under nutrient replete conditions, sessile thecate cells differentiate into motile slow swimmers that can proliferate as single cells or chains of cells connected through intercellular bridges^6,23,24^. In response to specific bacterial^25–27^ and algal^28^ cues, slow swimmers develop into rosettes, which are multicellular colonies that form through serial cell division^29^. As rosettes or slow swimmers starve, they become fast swimmers, and under prolonged nutrient limitation, fast swimmers differentiate into thecates that can proliferate under similarly harsh conditions^6^. The recapitulation of these cell type transitions in the laboratory make *S. rosetta* a promising model to investigate how a marine microeukaryote integrates environmental signals to produce distinct cell types. However, the environmental functions that *S. rosetta* cell types could perform remain to be uncovered, as choanoflagellate cell types have mostly been defined through their morphological features^3,6,29,30^ rather than molecular functions.

We refined previous *S. rosetta* transcriptomes^31^ to uncover functional differences between cell types reflected in their gene expression profiles. To improve those transcriptomes, we took advantage of methods to stably culture thecate, slow swimmer, rosette, and fast swimmer cell types in monoxenic strains of *S. rosetta* that feed on the bacterium *Echinicola pacifica*^23,24,32^. We also developed a method to preferentially lyse *S. rosetta* while discarding their feeder bacteria to harvest mRNA from *S. rosetta* with a single round of poly-A selection (Fig. S1) rather than previous methods that relied on multiple rounds of selection to deplete contaminating RNAs from bacteria^1,10^. A principal component analysis of these improved transcriptomes emphasized the difference between the thecate cell type compared to slow swimmers, rosettes, and fast swimmers (Fig. 1B) with the first principal component accounting for 86% of the variance and clearly separating thecates from all other cell types. While these results are consistent with previous gene expression profiles that found thecates separated from other types of cells^31,34^, the replicate measures in our data do not show a clear difference between slow swimmers, rosettes, and fast swimmers, suggesting that these three cell types are phenotypically plastic states.

**Figure 1:**
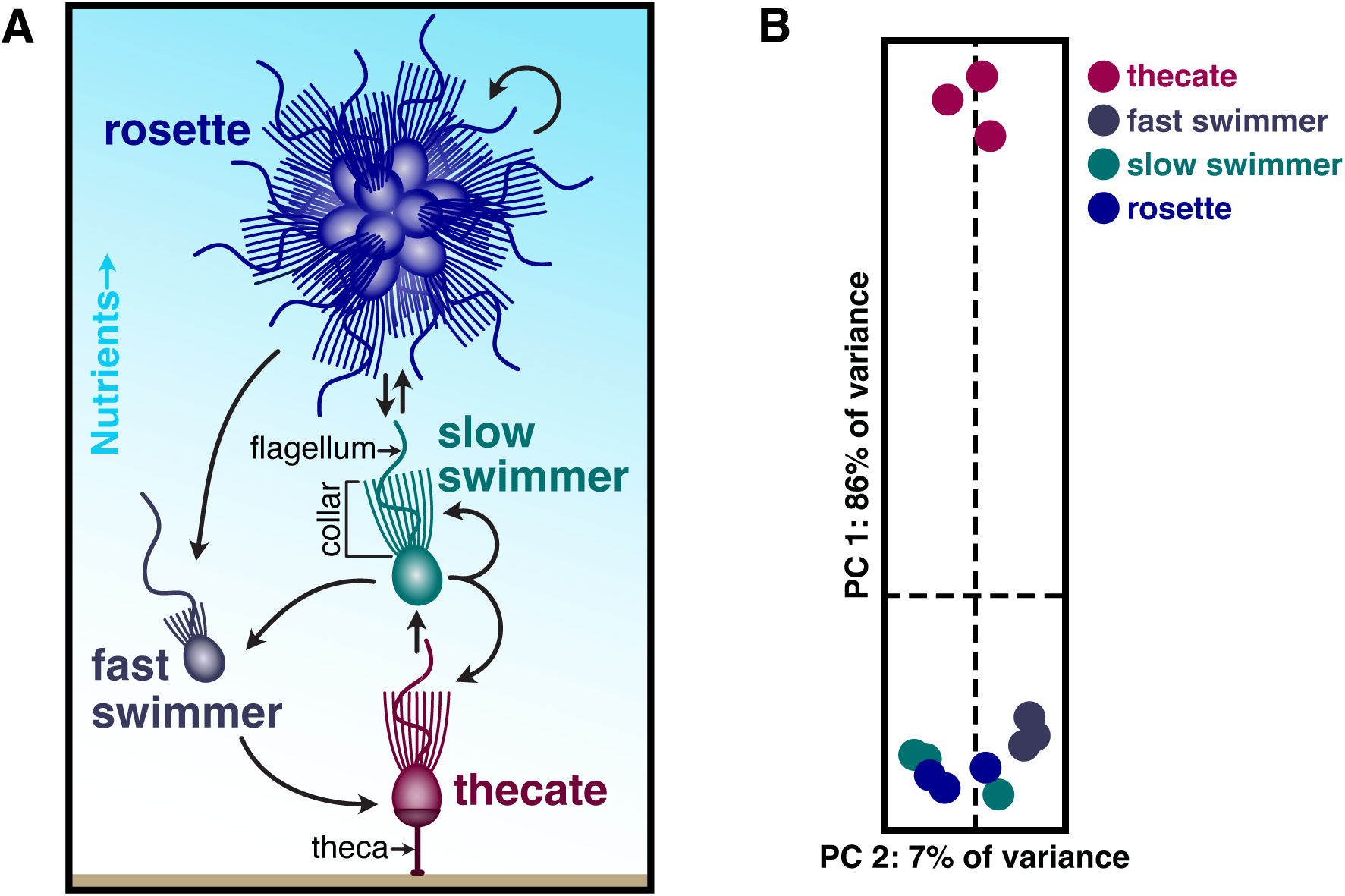
Thecates from the choanoflagellate *S. rosetta* are a distinct cell type. **(A)** *S. rosetta* differentiates into morphologically distinct cell types. This schematic shows four of the cell types from *S. rosetta* that were stably cultured to produce transcriptome profiles. All of these cell types display the common choanoflagellate cell architecture in which an apical flagellum is encircled by a collar of actin-filled microvilli (indicated on the slow swimmer) that enables choanoflagellates to phagocytose bacteria and particulate matter. When nutrients are abundant, slow swimmers (green) respond to bacterial and algal cues to develop into multicellular rosettes (blue) that form by serial cell division. Cultures of slow swimmers also form chains of cells through serial cell divisions, but those chains are easily disrupted by mechanical force. Under starvation, slow swimmers and rosettes become fast swimmers (grey), which have a reduced cell body and collar with a longer flagellum. Under sustained nutrient deprivation, fast swimmers differentiate into thecates (red), a type of cell that adheres to substrates by constructing an extracellular apparatus called a theca. Thecates still proliferate in low nutrient conditions by dividing into swimmers that build their own theca. By taking the supernatant of thecate cultures in high nutrient conditions, thecates can then differentiate into slow swimmers. **(B)** The transcriptome profile of thecates stands apart from all other cell types. A principal component analysis of triplicate RNA-seq profiles from each cell type (panel A) shows that the 86% of the variance between samples is attributed to the thecate transcriptome profile. All other cell types cluster closely together.

The contrast between thecates and other cell types prompted us to investigate what unique functions thecates may perform. By comparing thecates to slow swimmers, we identified a set of genes that reproducibly (*q* < 0.01, note that *q* is a *P*-value adjusted for multiple comparisons) displayed a greater than two-fold change in gene expression in thecates. We analyzed this set of genes for common gene ontologies (GO)^35–39^ that indicated functional categories enriched in thecates, keeping in mind that GO analysis may underestimate functional differences between cell types due to incomplete genome annotations in this emerging model system^40^. Genes upregulated in thecates were clearly enriched for functional modules (Fig. S2) implicated in cell signaling (e.g. nucleotide binding, protein kinase, protein phosphatase, calcium binding, metal ion transporter, guanylate cyclase and G-protein coupled receptor), gene regulation (e.g. transcription factor and ribosome component), and metabolism (e.g. nucleotide binding, oxidoreductase, glycosyl hydrolase, metal ion transporter, carbohydrate binding, sulfuric ester hydrolase, and ascorbic acid binding). In contrast, swimmers expressed modules for cell division, RNA processing, and oxidative metabolism (Fig. S3). This comparative analysis highlighted a gene (PTSG_09715) with 479-fold higher expression in thecates compared to slow swimmers (*q* < 10^-40^) that was annotated as an oxidoreductase (GO:0016491) composed of a single cytochrome b561 iron reductase domain (Pfam 03188). Given the role that oxidoreductases perform in diverse eukaryotes to regulate iron acquisition^41–44^, the strong induction of PTSG_09715 (henceforth called *cytb561a*) led us to hypothesize that thecates may have a different capacity to acquire iron from the environment compared to other cell types.

### *cytb561a* expression improves thecate cell growth with ferric colloids

Of the 2,820 genes that reproducibly displayed a two-fold or greater change in mRNA expression in thecates compared to slow swimmers, the change in *cytb561a* expression is in the ninety-eighth percentile (Fig. 2A). This large change in *cytb561a* expression may have been coupled to the thecate cell type program, may have been due to differences in strains to generate thecate and slow swimmer transcriptomes, or may have been an independent response to external nutrients that differed between the media for culturing slow swimmers and thecates for the transcriptomes (see methods). To distinguish among these possibilities, we measured *cytb561a* mRNA expression in slow swimmers and thecates from the same strain of *S. rosetta* after standardizing a culturing regime (Fig. 2B and Table S1). In this protocol, slow swimmers and thecates were grown in the same media formulation and then depleted of iron. Afterwards, we reintroduced iron either as soluble ferric EDTA (Fe^3+^•EDTA) or as insoluble ferric colloids (Fe^3+^(oxyhydr)oxides)^40^. Both forms of iron are ecologically important, for models based on environmental sampling predict that 40% of the world’s surface ocean waters rely on colloidal sources of iron and 18% on soluble iron chelates^11^. With either source of iron, the expression of *cytb561a* mRNA increased in thecates compared to slow swimmers (Fig. 2C and S4 A-C): 89-fold with ferric EDTA and 66.2-fold with ferric colloids (two-way ANOVA, *P* < 0.005). This result supports the conclusion that thecates are a stable cell type that expresses *cytb561a* as part of its differentiation regulon.

**Figure 2:**
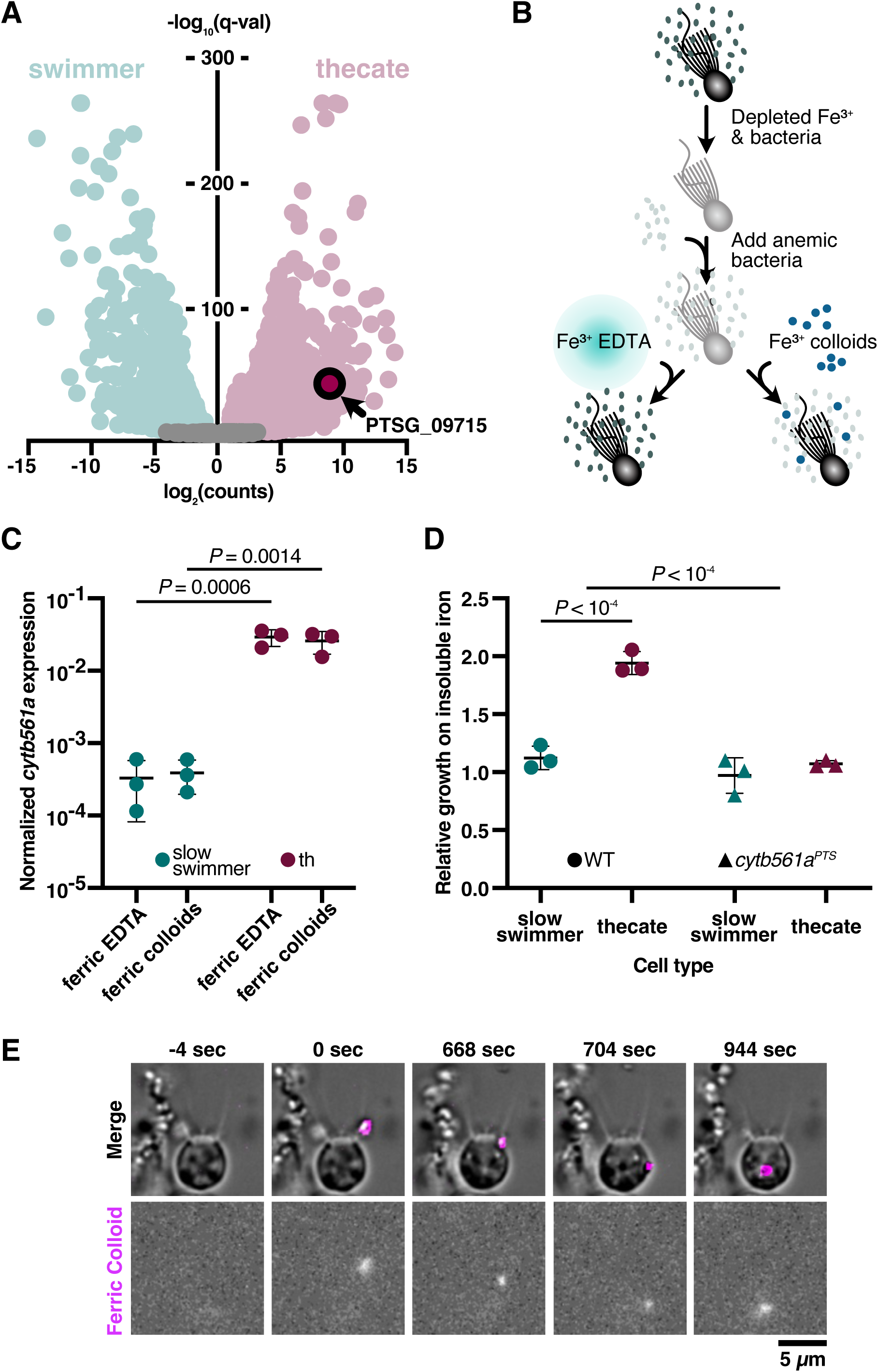
Thecates utilize ferric colloids through the expression of an iron reductase, *cytb561a*. **(A)** Thecates highly express an iron reductase. The log_2_ transformed ratio of transcript abundances (counts) from thecates and slow swimmers (x-axis) plotted against *q*-values (y-axis) highlights transcriptome changes in each cell type.. Among the genes with transcript abundances that reliably (*q* < 0.01) changed more than two-fold in thecates (red) compared to swimmers (green), PTSG_09715 is in the 98^th^ percentile. This gene, which is annotated as an iron reductase, encodes a single domain: cytochrome b561 (Pfam 03188). **(B)** A method to swap iron sources that support choanoflagellate growth. In preparation for iron limitation assays, *S. rosetta* was passaged in low-nutrient media and depleted of iron. Cells were then washed multiple times and inoculated with iron-limited feeder bacteria. At the same time, ferric EDTA (Fe^3+^•EDTA) or ferric colloids (Fe^3+^) were provided. Cultures grew for 48 hours at 27°C before assaying *cytb561a* expression (Fig. 2C and S2) and growth (Figs. 2D and S4) **(C)** *cytb561a* expression is part of the thecate regulon, and not determined by external iron conditions. The expression of *cytb561a* was monitored by RT-qPCR in cultures of slow swimmers (green) or thecates (red) that were grown with either ferric EDTA or ferric colloids. The expression of *cytb561a* was normalized to *cofilin* (PTSG_01554), a eukaryotic gene that displays high, consistent expression across all *S. rosetta* cell types. Independent triplicates were performed and *P-*values were calculated from a two-way ANOVA. **(D)** Thecates require *cytb561a* for increased proliferation with ferric colloids. To account for differences in growth between slow swimmers (green) and thecates (red), the cell density of cultures grown with ferric colloids was normalized to cultures grown with ferric EDTA. With this metric, a ratio greater than one indicates that the cell type displays increased growth with ferric colloids; whereas, a ratio less than one indicates the converse. A premature termination sequence introduced at position 151 in *cytb561a* with CRISPR/Cas9 genome editing produced a mutant allele (*cytb561a^PTS^*) with a ten-fold reduction in *cytb561a* mRNA levels and stop codons that would truncate the protein translated from this transcript (Fig. S2 and Table 2). Unlike the wild-type strain (circles), thecate cells with *cytb561a^PTS^*displayed no improved growth with ferric colloids (triangles). **(E)** Thecates ingest ferric colloids through phagocytosis. Time courses of wild-type thecates incubated with fluorescently labeled ferric colloids (see methods on ferric colloid labeling). Fluorescent ferric colloid particles were tracked (magenta outline) and observed being ingested and internalized through phagocytosis. The zero time point indicates initial contact with the ferric colloid particle.

To determine the functional consequence of increased *cytb561a* expression in thecates, we compared the growth of thecates and slow swimmers in the presence of 100 µM ferric EDTA or ferric colloids (Fig. 2D). Thecates grew to a two-fold higher cell density in the presence of ferric colloids (two-way ANOVA, *P* < 10^-4^), yet slow swimmers showed no change (two-way ANOVA, *P* > 0.1). This effect was observed at lower concentrations of iron (50 µM), but higher concentrations of iron (≥200 µM), favored greater cell proliferation in ferric EDTA (Fig. S5A). Thecates also grew better than slow swimmers with other ferric chelates, such as ferric EGTA and ferric pyoverdine (Fig. S5B). Importantly, the differences in *S. rosetta* growth were unlikely due to indirect effects from the prey bacteria, for *E. pacifica* grew similarly in all iron conditions, likely due to the small concentration of iron (3.43 ± 1.01 µM) that dissolved from the ferric colloids (Fig. S5 C-E).

The increased proliferation of thecates in the presence of ferric colloids did depend on the expression of *cytb561a*. With an improved genome editing pipeline (see methods and Fig. S6), we altered *cytb561a* expression by introducing a premature termination sequence^46^ at position 151 of *cytb561a* to incorporate missense mutations and a polyadenylation signal (Fig. S7), which diminished mRNA expression in thecates by 3.6-fold compared to wild-type (Fig. S4D, one-way ANOVA, *P* = 0.004). This mutant allele, *cytb561a^PTS^*, eliminated the difference in cell proliferation between slow swimmers and thecates (Fig. 2D, two-way ANOVA *P* > 0.1). A comparison of *cytb561a^PTS^* and wild-type growth dynamics (Fig. S8) showed that the enhancement of thecate growth with ferric colloids was primarily due to a two-fold larger carrying capacity and a shorter doubling time (17.9 h vs 13.2 h, two-way ANOVA, *P* = 0.035). To derive iron from ferric colloids for enhanced cell growth, thecates can directly ingest iron particles, as seen in time-lapse images of thecates feeding on ferric colloids embedded with a fluorescent polysaccharide (Figs. 2E & S9). Taken together, these results suggest that the increased expression of *cytb561a* in thecates confers a cell-type-specific function to assimilate iron from insoluble ferric colloids for faster growth to higher cell densities.

### Eukaryotes have broadly conserved a core iron acquisition toolkit

Although *cytb561a* expression was necessary for the increased proliferation of thecates when grown with ferric colloids, we were uncertain how this iron reductase factored into the pathway(s) for iron acquisition in *S. rosetta*, as iron transport pathways have not previously been examined in choanoflagellates. Therefore, we compiled an inventory of iron import and export proteins that have been characterized in diverse model eukaryotes (e.g. humans^44^, *S. cerevisiae*^47^, *P. tricornutum*^41^*, C. reinhardtii*^43^, Fig. S10) to survey the distribution of those proteins across Holozoa (the taxonomic group that includes Animalia, Choanoflagellata, Filasterea, and Teretosporea) and a curated set of eukaryotes^48^. We found that nearly all of the proteins for iron import, export, reduction, and oxidation are widely distributed among eukaryotes, suggesting a deep ancestry of iron acquisition pathways from the last common eukaryotic ancestor (Fig. 3A and S11).

**Figure 3:**
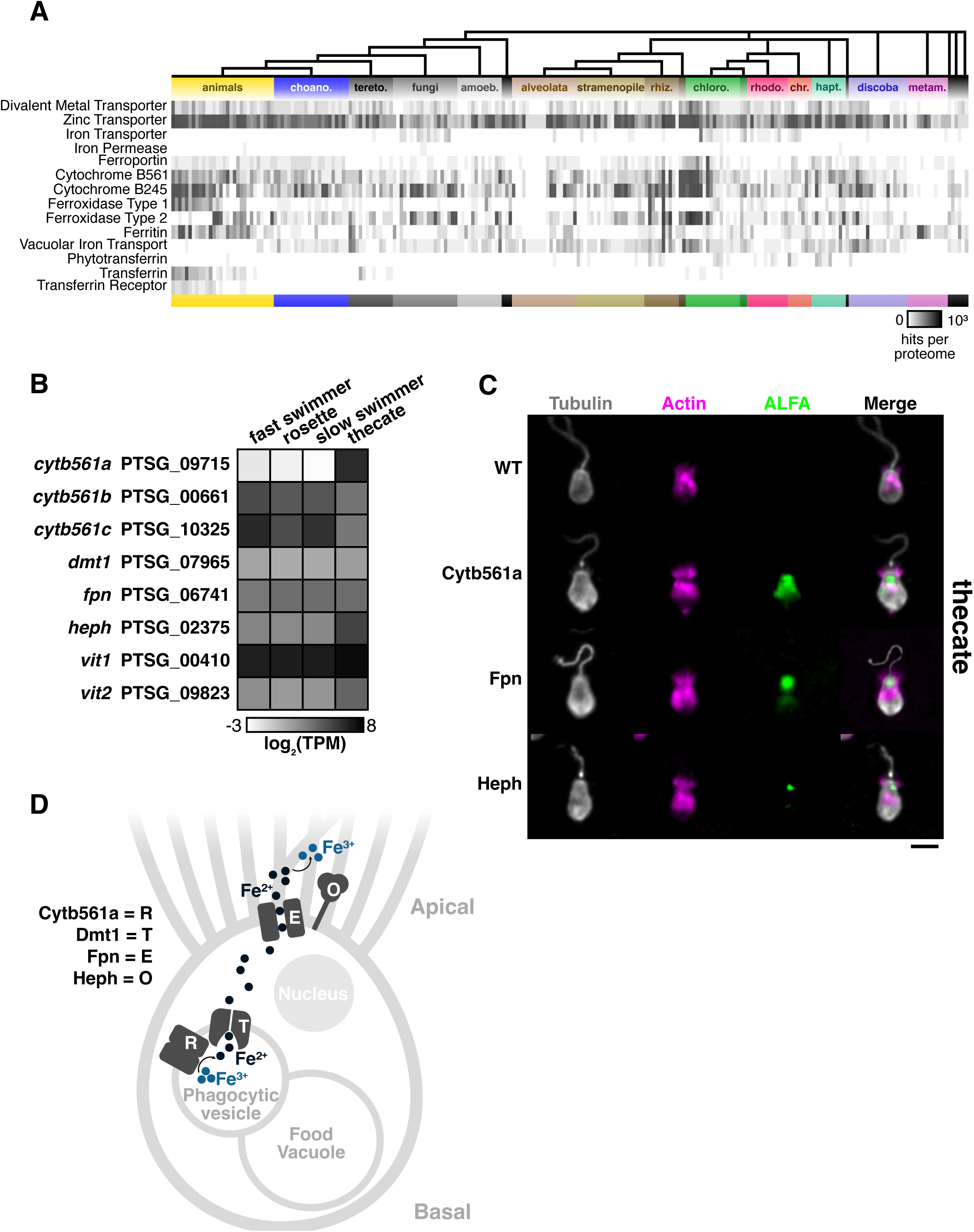
Iron acquisition pathways in choanoflagellates and animals evolved from a toolkit widely conserved in eukaryotes. **(A)** Distribution of iron acquisition proteins across diverse eukaryotes. Iron acquisition proteins characterized in model eukaryotes were compiled into a list to search a curated database of proteomes predicted from the genomes and/or transcriptomes of diverse eukaryotes. The iron acquisition proteins are sorted into these categories from the top to bottom rows: transporters, reductases, oxidases, iron storage, and sharing/binding. Each column indicates a single species, and major eukaryotic lineages are denoted by color. The number of unique protein hits identified in each species is shown in greyscale. **(B)** Cell-type expression of iron acquisition genes identified *S. rosetta* cell types. Genes are denoted by unique identifiers for *S. rosetta* genes and gene names that were given in this work based on their homology with animals. Expression values are averaged triplicate values of transcripts per million (TPM) from cell type transcriptomes (Fig. 1). Of all genes and paralogs, *cytb561a* exhibits the most striking differential regulation. **(C)** The localization of iron acquisition proteins in thecates. An ALFA epitope was engineered into endogenous genetic loci at the carboxy terminus of the protein coding sequences. Although this tag was engineered into all iron acquisition genes (except the vacuolar transporters), only the proteins displayed in this panel were visible in western blots (Fig. S7). Immunofluorescent staining of ALFA-tags, tubulin and phalloidin indicate the localization of tagged protein relative to cytoskeletal markers of cellular architecture. Images of individual cells are representative of more than three independent staining experiments in which six or more images were captured of fields containing more than ten cells. Scale bar is 5 µm. **(D)** A putative iron acquisition pathway in *S. rosetta*. Iron is internalized by phagocytic vesicles derived at the base of the collar. Inside, Cytb561a reduces iron for transport into the cytosol by Dmt1. Iron is then assimilated by the cell. To maintain iron homeostasis, Fpn exports iron out of the cell, and Heph aids in maintaining a favorable concentration gradient for export.

Our searches also revealed notable evolutionary patterns that correspond to the unique biology of certain eukaryotic lineages. For example, we found significant losses of iron toolkit genes in some lineages. Apicomplexa, a taxon that includes the intracellular parasites *Cryptosporidium parvum*, *Toxoplasma gondii*, and *Plasmodium falciparum* has a reduced set of iron acquisition genes, consistent with the origin of *de novo* iron acquisition pathways for siphoning iron from their hosts^49^. Moreover, we observed that some iron utilization genes are more taxonomically restricted than others: Genes first characterized in ascomycete yeasts were either primarily found in Fungi and Archaeplastida (e.g. iron transporter^50^) or only present in a few eukaryotes (e.g. iron permease^51^). Similarly, transferrin and phytotransferrin were primarily found in Holozoa and Archaeplastida, respectively, as previously shown^52^. Notably, the presence of transferrin in Animalia and Teretosporea indicated that transferrin was present in the last common ancestor of Holozoa, yet the transferrin receptor that facilitates the cellular internalization of iron-bound transferrin appears to have originated in Deuterostomes (Fig. S12), highlighting the origin of a mechanism to share iron between cells in the multicellular bodies of animals. Finally, we also observed that the presence of genes encoding vacuolar iron transporters and ferritin, which are both involved in iron storage, was slightly anticorrelated (*r* = −0.380, *P* < 10^-8^). Overall, this catalog of iron utilization proteins indicates that eukaryotes have variably retained, amplified, or lost genes from an ancestral toolkit for the acquisition, storage, and export of iron.

Choanoflagellates and animals appeared to share a core pathway for acquiring iron from their external environment. Iron acquisition in animals has most extensively been studied in vertebrate gut epithelia^53^. The apical surface of gut epithelial cells absorbs dietary iron by reducing ferric cations to ferrous cations through the activity of duodenal cytochrome B561 (DCYTB) before the divalent metal ion transporter (DMT1) brings ferrous cations inside of cells. On the basal surface of gut epithelia, ferroportin (FPN) secretes ferrous cations into the bloodstream, where they are oxidized to ferric cations near the membrane by hephaestin (HEPH). In our catalog of iron utilization genes, all choanoflagellates possessed homologs of *DCYTB*, *DMT1*, and *FPN*, but *HEPH* (Fig. 3A, row 8–ferroxidase type1) was only present in *S. rosetta* and 9 other species (of 22 surveyed^54^) – perhaps due to the lack of complete genomes for many of those species. Differences between choanoflagellates and animals were evident in iron storage mechanisms, as choanoflagellates mostly possessed vacuolar iron transport proteins, which have been studied in *S. cerevisiae*^55,56^; whereas, animals broadly conserved ferritin. As we searched for homologs of iron acquisition genes in *S. rosetta*, we found single copies of *dmt1*, *fpn*, and *heph* and identified two vacuolar iron transporters (*vit1* and *vit2*). Three cytochrome b561 genes were present, yet among all the conserved iron utilization genes, only *cytb561a* exhibited strong differential expression between cell types (Fig. 3B), emphasizing its cell-type-specific role in thecates.

Because the apico-basal localization of iron acquisition genes facilitates the unidirectional import of iron in animal epithelia^53^, we tested if iron transport in *S. rosetta* shared a similar architecture (Fig. 3C). To do so, we used genome editing at endogenous loci to introduce an epitope tag (ALFA-tag^57^) in all the iron utilization genes (Fig. S7). Only Cytb561a, Fpn, and Heph were visible in western blots of thecates (Fig. S13A), so we focused on those proteins to visualize by immunofluorescence. In thecates, Cytb561a localized intracellularly toward the apical half of the cell but not in any pattern resembling organelles^53^. Cytb561a was undetectable in slow swimmers (Fig. S13B), in contrast to thecates that stably expressed Cytb561a independent of iron availability (Fig. S14), both of which are consistent with mRNA expression levels (Fig. 2C). In both slow swimmers and thecates, Fpn and Heph localized near the apical membrane inside of the collar (Fig. S15). The similar co-localization of Fpn and Heph would be consistent with their coordinated function as an efflux pump and oxidase as described in animals^53^. Importantly, the basal localization of these proteins appears to be a feature of animal epithelia that is not shared with choanoflagellates and may have evolved alongside the mechanisms to establish and maintain cell polarity^59^.

Based on the functions of homologous genes and the localization of Cytb561a, Fpn, and Heph, we propose a model for iron acquisition in *S. rosetta* thecates that may extend to other choanoflagellates (Fig. 3D). After internalizing ferric colloids by phagocytosis, Cytb561a reduces ferric cations in endosomes where we surmise that Dmt1 transports ferrous cations into cells – although Dmt1 was undetectable by immunohistochemistry (Fig. S14). As slow swimmers do not express Cytb561a, *S. rosetta* may acquire iron directly from bacterial prey digested in food vacuoles or rely on Cytb561b or Cytb561c to reduce ferric cations. In both slow swimmers and thecates, we hypothesize Fpn exports ferrous cations at the apical surface where Heph can oxidize it.

### Cytb561 paralogs display distinct biochemical properties and global distributions

The variation in the number of cytochrome b561 paralogs across eukaryotes led us to examine the evolution and function of the three cytochrome b561 paralogs that we identified in *S. rosetta*. While the expression of *cytb561a* in thecates was clearly necessary for improved growth in the presence of colloidal iron, slow swimmers still grew in those conditions as did *cytb561a^PTS^*thecates (Fig. 2D). These observations and the steady mRNA expression of *cytb561b* and *cytb561c* across cell types (Fig. 3C) would be consistent with these paralogs sustaining a baseline ability to acquire iron, even if we could not detect their protein products by immunohistochemistry (Fig. S13). The potential for functional redundancy among these paralogs also led us to consider if choanoflagellates and other eukaryotes display lineage-specific expansions of this gene family that may have resulted in differing capacities to acquire iron.

To better compare the cytochrome b561 paralogs in *S. rosetta*, we first built a phylogenetic tree^60^ from an alignment^61^ of Cytb561 proteins from Amorphea (Fig. 4A), a eukaryotic supergroup that includes Holozoa and Amoebae^48,62^. The unrooted tree of cytochrome b561 proteins robustly supported (ultrafast bootstrap^63^ > 88) three distinct groups. Groups B and C contained protein sequences from each major clade in Amorphea and its sister group CRuMs (Collodictyonida, Rigifilida and Mantamonadida). Strikingly, sequences from Holozoa dominated Group A, as a single protein from Amoebozoa was the only sequence from outside of Holozoa that fell into this group. Notably, Group A not only contained mammalian DCYTB proteins but also Cytb561a from *S. rosetta*, supporting the annotation of this gene as an DCYTB ortholog. The Cytb561b and Cytb561c proteins from *S. rosetta* fell into Groups B and C, respectively.

**Figure 4:**
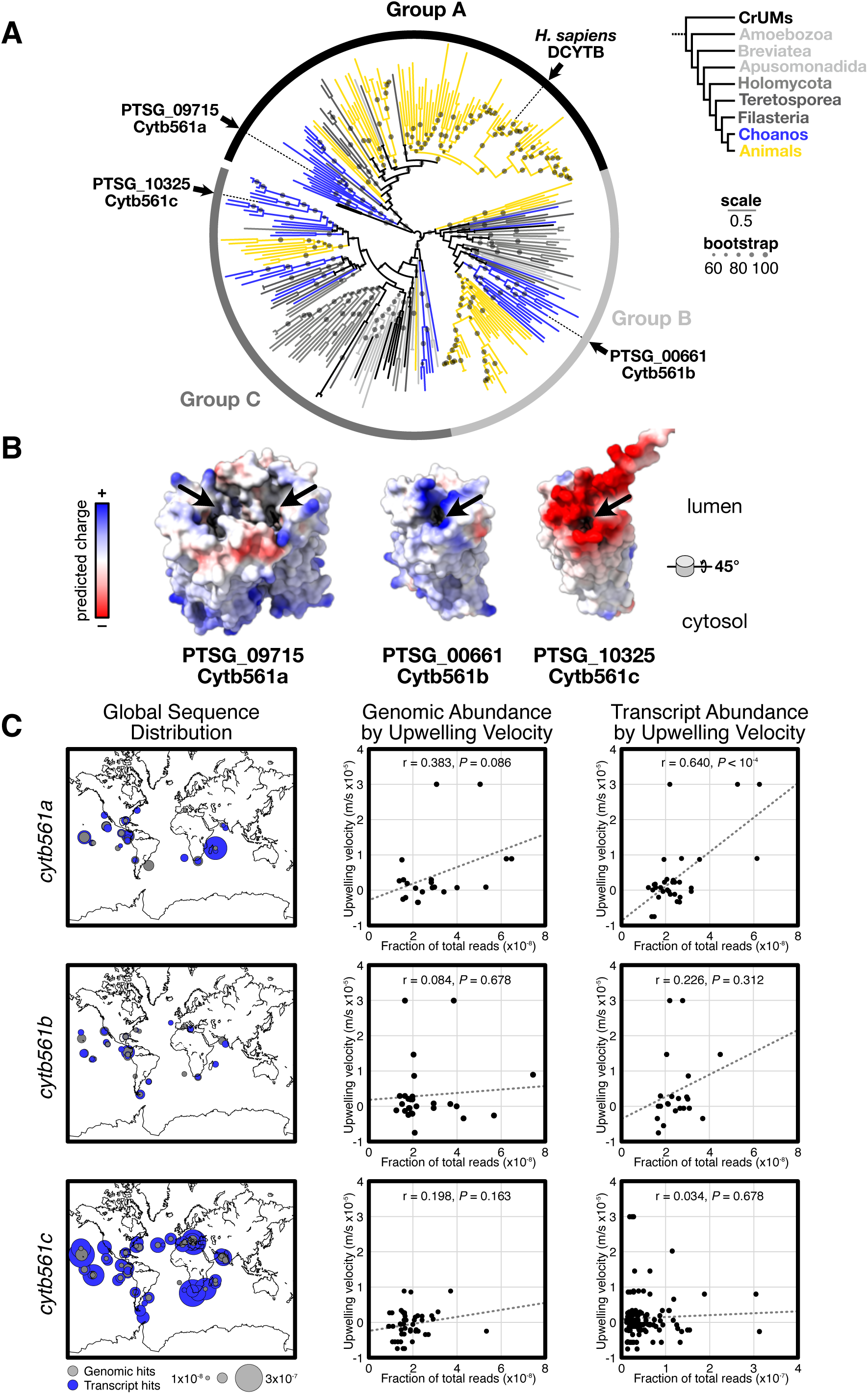
Cytb561 paralogs possess distinct biochemical properties and appear at different oceanic locations. **(A)** Phylogeny of iron reductases reveals Cytb561a is an ortholog of animal DCYTB. Each paralog of *S. rosetta* falls into distinct clades of cytochrome b561. Circles on each branch are proportional to bootstrap values greater than sixty. Scale bar indicates the average number of substitutions per site. Branch colors correspond to clades as shown in the legend: choanoflagellata (choano.), tereto. Teretosporea (tereto.), amoeba (amoeb.), rhizari (rhiz.), rhodophyta (rhodo.), chromista (chr.), haptophyta (hapt.), metamonoda (metam.) **(B)** Predicted structures of *S. rosetta* Cytb561 paralogs show differences in dimerization and substrate-binding interfaces. Alphafold (v3) predictions show Cytb561a as the only paralog forming homodimers. Predicted electrostatic surfaces Cytb561 paralogs show different charge distributions around the substrate binding pocket (arrows). Models are angled 45° for viewing the lumenal surface where iron binds. **(C)** *cytb561a* transcript abundance correlates with ocean upwelling velocities. Maps (left panels) show the genomic (grey) and transcript (blue) abundance of each paralog. Plots show genomic (middle panels) and transcript (right panels) abundance on the y-axis and average upwelling velocities on the x-axis. Note, the x-axis for *cytb561c* transcript abundance is scaled differently than for *cytb561a* and *cytb561b*. *r* indicates Pearson correlation and *P-*values were calculated from a t-test.

Because each of the *S. rosetta* paralogs was part of a different group of cytochrome b561 proteins, we examined^64^ multiple sequence alignments (Figs. S16,17,19) and their predicted structures^65^ (Fig. S18) to evaluate if the deep evolutionary divergence was reflected in their biochemical characteristics (Fig. 4B), using the crystal structure of human DCYTB as a reference^66^. DCYTB forms a homodimeric complex with each monomer folding into six transmembrane helices that sandwich two heme groups in the interior of the protein. These heme groups mediate the transfer of an electron from an ascorbate molecule on the cytosolic side of the protein to ferric cations on the lumenal side. Thus, in our comparisons of multiple sequence alignments, we focused on the residues in DCYTB that bind ascorbate, mediate homodimerization, and attract iron. First, all homologs appear to have conserved positively charged residues that form the pocket to bind negatively charged ascorbate^66^ (Fig. S16).

Second, only proteins from Group A conserve residues that mediate dimerization (Fig. S17), which was also reflected in predicted dimeric structures of paralogs from *S. rosetta* (Fig. S18). Third, among the Cytochrome b561 groups, we noticed significant differences on the lumenal surface that forms the binding site for ferric cations (Figs. 4B, arrow and S19). Homologs within Group A, which contains Cytb561a and DCYTB, possess a mix of positive and negative charges, indicating Group A paralogs can bind iron and coordinated ligands^66^. Group C homologs have largely anionic surface charges at the putative iron binding site, which could plausibly bind iron as well. However, the positively-charged binding pockets conserved across Group B homologs may repel rather than bind iron cations. Notably, the differences in charge distribution among Cytochrome b561 groups appear to be general features of those groups (Fig. S19C), yet electrostatic properties can diverge within those lineages (Fig. S19D). With each paralog from *S. rosetta* possessing distinct biochemical properties near the iron binding site, we hypothesize that the improvement of thecate growth with ferric colloids may not only be due to the mass action from more Cytb561 proteins but also through the distinct biochemical activity of Cytb561a.

If the biochemical activity and cell-type-specific expression of Cytb561 paralogs enable choanoflagellates to acquire iron from different environmental sources, then we expect that the expression of each paralog may be enriched in different ecological settings. To test this hypothesis, we searched for each paralog in an open repository (Ocean Gene Atlas^8,9^) of metagenomes and metatranscriptomes from marine environments. When we visualized the distribution of each paralog, we found that they varied in both location and transcript abundance but were similarly present in metagenomic reads (Fig. 4C) The most abundant and cosmopolitan paralog was *cytb561c*. Although present at most of the sites, the transcript abundance of *cytb561b* was ∼10-fold lower than *cytb561c*. The abundance of *cytb561a* had a similarly low abundance as *cytb561b,* yet its distinct presence near coastlines and the equator resembled the locations of marine upwellings^67^ (Fig. S20A).

Importantly, upwellings are a major source of remineralized iron colloids rising from ocean depths^68,69^, and upwelling velocities may serve as a better proxy for the presence of colloidal iron than predictions of soluble iron concentrations^10^. Therefore, we correlated the abundance of the genomic and transcript reads for each paralog to average vertical velocities at a depth of 100 meters for each sample location (Fig. 4C, S20B). Of all the comparisons, the only robust correlation was between *cytb561a* transcript abundance and vertical velocity (*r* = 0.64, *P* < 10^-4^), as the transcript abundances of the other paralogs exhibited only weak positive correlations that were probably random (*P* > 0.3). The positive correlation between *cytb561a* transcripts and upwelling velocities indicate ecological contexts where choanoflagellates may be acquiring insoluble iron colloids for their growth. We cannot, however, determine if the choanoflagellates in those regions are thecates expressing *cytb561a* as in *S. rosetta*, or choanoflagellates with a different regulatory mechanism that evolved uncoupled *cytb561a* expression from thecate development. Nonetheless, these observations connect *cytb561a* expression to an environmental feature and places choanoflagellate biology in a global ecological framework.

Iron is an essential micronutrient affecting the growth and productivity of phytoplankton^70^. Field measurements and ocean scale models have found that iron particulates are the major source of iron for 40% of the world’s surface ocean waters^11^. In addition to photolytic reduction^14,71^ and bacterial siderophores^72,73^, the ingestion of iron particles by phagotrophic protists is a major mechanism to solubilize iron and can alleviate iron limitation for cell-walled organisms that rely on obtaining iron in soluble forms^13–15^. As efficient phagotrophs^74,75,4,76^, choanoflagellates ingest iron particulates (Figs. 2E and S9), yet iron assimilation from ferric colloids depends on the expression of a specific Cytb561 paralog in only one cell type of *S. rosetta*. This dependency emphasizes the importance of integratively factoring protein evolution^77^ and gene expression during life history transitions^1,2^ in models for determining^78^ the ecological roles of microeukaryotes in the acquisition and distribution of limiting nutrients throughout their surrounding environment. Moreover, the synthesis of cytological, genomic, and ecological data may help to interpret how the rise in oxygen that resulted in the depletion of soluble ferrous cations from the oceans influenced eukaryotic evolution^77,79,80^, including the emergence of animals^81^.

The cell-type-specific expression of *cytb561a* not only illustrates how cellular differentiation impacts the environmental function of a choanoflagellate but also informs how cellular differentiation evolved as a defining feature of animal multicellularity^16,19,17,82,20,21^. Choanoflagellate cell types have been primarily defined by their morphology and motility^4,6^. In addition to those cell biological characteristics, we show here that thecates have a distinct ability to utilize ferric colloids. The coupling of this function to cell differentiation stands in contrast to the regulation of iron utilization in other microeukaryotes that transiently regulate iron utilization genes in response to external iron concentrations^56,83,84^. The functional difference between slow swimmers and thecates more closely resembles the stable differentiation of cell types during animal development^85–87^. Taken together, these observations support models for the origin of animal cell differentiation in which the last common holozoan ancestor differentiated into unique cell types at each life history stage^88,16,17,20,22^. Evolution along the animal stem lineage coupled cell differentiation to multicellular development, resulting in multicellular bodies composed of functionally distinct cells^89^. Importantly, mechanisms to distribute resources from cells that acquire nutrients from the external environment (e.g. epithelial cells^90^) to cells that reside internally helped to transform the functions of various cell types into a common organismal physiology^18^. As the evolution of transferrin and its receptor illuminates (Figs. 3A and S12), the multicellular physiology of animals has likely been cobbled together through the co-option of mechanisms for nutrient acquisition and cellular trafficking.

## Materials and Methods

### Culturing S. rosetta

To grow cell types of *S. rosetta* for the generation of transcriptomes, slow swimmer and rosettes were grown in 5% Sea Water Complete (SWC) Media^23,91^ (Table S1), seeded at 10^4^ cells/ml and grown for 48 hours at 22°C. Rosettes were induced with outer membrane vesicles (OMVs) harvested from the bacterium *Algoriphagus machipongonensis*^26^. Fast swimmers were prepared the same way as slow swimmers, but were grown for 72 hours at 22°C and transitioned to 30°C for 2 hours and 45 minutes. Thecates were grown from the strain HD1, a strain which is constitutively in the thecate cell type, in 10% Cereal Grass Media 3 (CGM3)^23,91^ (Table S1), seeded at 10^4^ cells/ml and grown for 48 hours at 22°C.

For iron acquisition experiments and standard culturing of *S. rosetta*, all cell types were grown in 25% Red Algae (RA) Media^92^ (Table S1) with the feeder bacteria *Echinicola pacifica*. Cultures were grown at 18°C for maintaining cultures and shifted to 27°C one day prior to conducting iron acquisition experiments.

### RNA extraction

**Protocol Link:** dx.doi.org/10.17504/protocols.io.261ge4bojv47/v1

In a previous paper^91^, we developed a lysis buffer to extract RNA from *S. rosetta* that has subsequently been used in other work for RNA^30,93,94^ and protein^95^ extraction. Here we show that this buffer was developed to preferentially lyse *S. rosetta* in cultures that have an abundance of the prey bacterium *E. pacifica*. To preferentially lyse *S. rosetta*, we reasoned that sterol-based detergents, such as digitonin, would more effectively disrupt membranes with sterols, like those of *S. rosetta* and other eukaryotes^96^ rather than bacterial membranes that largely lack the sterols^97^. Additionally, the lysis buffer contains RNase inhibitors (RNaseIN and heparin) and translation inhibitors to decrease the degradation of mRNA. We evaluated the efficacy of this buffer by tracking ribosomal RNA from *S. rosetta* and *E. pacifica* in lysed samples that were centrifuged to separate the supernatant and pellet, in which we found that *S. rosetta* ribosomal RNAs were enriched in the supernatant and bacterial ribosomal RNAs were enriched in the pellet (Fig. S1). The optimization of this procedure resulted in the following method to obtain RNA samples from each cell type of *S. rosetta*:

After counting the cell concentration in a culture of *S. rosetta* feeding on *E. pacifica*, a volume yielding 10^7^ cells was centrifuged at 2600xg for 5 min at 4°C. The pellet from this sample was resuspended in 100 µl of preferential lysis buffer (20 mM Tris-HCl, pH 8.0; 150 mM KCl; 5 mM MgCl_2_; 250 mM Sucrose; 1 mM DTT; 10 mM Digitonin; 1 mg/ml Sodium Heparin; 1 mM Pefabloc SC; 100 µg/ml Cycloheximide; 1 tablet/ 5 ml buffer EDTA-free Protease Inhibitor Tablet [Sigma-Aldrich Cat. No. 11836170001]; 0.5 U/µl Turbo DNase [Thermo Fisher Scientific, Cat. No. AM2239]; 1 U/µl SUPERaseIN [Thermo Fisher Scientific, Cat. No. AM2696]) and incubated on ice for 10 min. Cells were triturated through a 30G needle ten times. Afterwards, the insoluble debris was pelleted by centrifugation at 6000xg for 10 min at 4°C. The supernatant was removed and adjusted to a volume of 100 µl with RNase-free water. Total RNA was purified on a silica membrane with the RNA clean-up protocol from the RNAeasy kit (Qiagen, Cat. No. 74104).

### RNA library preparation and sequencing

In preparation for sequencing, the quality of total RNA from *S. rosetta* samples was assessed by Agilent Bioanalyzer 2100 (Fig. S1C) with the Agilent RNA 6000 Nano Kit (Agilent, Cat. No. 5067-1511). Then total RNA (500 ng per sample) was purified by one round of polyA mRNA selection with oligo-dT magnetic beads (NEB, Cat. No. S1550S), converted to cDNA using KAPA mRNA HyperPrep kit (KAPA biosystems, Cat. No. KK8580), and indexed-adapter ligated with KAPA single-indexed adapter kit (KAPA biosystems, Cat. No. KK8701). Final library quality was determined by Bioanalyzer 2100 using the Agilent High Sensitivity DNA Kit (Agilent, Cat. No. 5067-4626) and pooled together after normalizing samples based on their quantity from qPCR. Sequencing was carried out by the QB3-Berkeley Genomics core (QB3 Genomics, UC Berkeley, Berkeley, CA, RRID:SCR_022170), on the Illumina HiSeq 4000. All samples were pooled and run on a single lane with 12.4 to 61.3 million reads per sample. Reads were demultiplexed, and checked for quality by fastqc (Babraham Bioinformatics). Metadata for sequencing can be found in Table S2.

### RNA-seq quantification and gene enrichment analysis

RNA-seq reads from each sample were mapped to a reference *S. rosetta* transcriptome (https://ftp.ensemblgenomes.ebi.ac.uk/pub/protists/release-58/fasta/protists_choanoflagellida1_collection/salpingoeca_rosetta_gca_000188695/cdna/) using kallisto^98^. Transcript abundance and differential expression were quantified with sleuth^99^. Gene enrichment for thecates was quantified by selecting genes with *q* < 0.01 and log_2_ (fold-change + 0.5) > 1. The resulting genes were assigned GO terms using DAVID Bioinformatics^37,38^ where *S. rosetta* gene IDs could be identified by “ENSEMBLE_GENE_ID”. For ease of interpretation, assigned GO terms were collapsed down using REVIGO^100^ and further assigned to functional categories in *S. rosetta* (TaxID: 946362) by GO Slim^39^ (Table S3).

### S. rosetta growth with different sources of iron

#### Media Preparation

To test the ability of *S. rosetta* cell types to differentially utilize iron sources, we developed a method to culture *S. rosetta* under iron limitation. We created an iron depleted media formulation named 4% Peptone Glycerol (PG) Media (Table S1). We added iron to this media from two different sources: Ferric EDTA was prepared by dissolving FeCl_3_ and EDTA to a final concentration of 1.46 mM in artificial seawater (ASW [Ricca Chemical Company, Cat. No. R8363000-20F]) and sterile filtering through a 0.22 µm polyethersulfone (PES) filter. Ferric colloids were prepared by adding a small volume of 70% (v/v) ethanol to FeCl_3_(*s*) to sterilize the solid; after evaporating the liquid in a biological safety cabinet, 1.46 mM FeCl_3_ was prepared in ASW. The solution was heated to 50°C for 10 minutes to precipitate ferric (oxyhydr)oxides, or ferric colloids^45^.

#### Fluorescent ferric colloid preparation

Ferric colloids were prepared as above, except during the 50°C incubation, a fluorescent dextran (10 kD anionic dextran conjugated to tetramethylrhodamin [Thermo Fisher, Cat. No. D1868]) was added to a final concentration of 20 µg/ml. This dextran co-precipitated with the ferric colloids, thereby embedding fluorophores within the colloids. The precipitated ferric colloids were pelleted by centrifugation at 4600 x g for 10 min at room temperature. The pellet was resuspended in ASW supplemented with 2% (v/v) dextranase (Sigma aldrich, Cat. No. D0443) to digest any excess dextran during a 10 min incubation at 60°C. After pelleting the ferric colloids again, they were resuspended to ∼1.46 mM in ASW. Finally, the resuspended colloids were ultrasonicated for two, 1 min cycles (pulsed for 1 sec on, 1 sec off) to create smaller particles for easier feeding and imaging, as larger particles broke down slowly and often obscured cells during imaging.

#### Determining iron concentration in media

To determine the background concentration of soluble iron in our newly formulated media and the amount of labile iron liberated from the ferric colloids, we used a modified protocol^13^ to measured soluble iron concentrations in 4% PG and 4% PG with 100 µM ferric colloids. To measure these iron concentrations, 50 ml of 4% PG media was incubated at 27°C for 6 hours with and without 100 µM ferric colloids. The media was then centrifuged at 4600xg for 15 minutes at room temperature. The top 45 ml of supernatant was taken for the following steps. To capture and concentrate the soluble iron in the supernatant, 125 µM 8-Hydroxyquinoline (oxine) was added to chelate labile iron, and 5 ml of 1M MES, pH 6 was added to aid in oxine’s solubility in the media and its binding to iron, which is facilitated by lower pH. These solutions were incubated for 24 hours protected from light, and then filtered through a Sep-Pak C18 column (Waters, Cat. No. WAT036800) to bind oxine-iron complexes. The column was then washed with 5 ml water and eluted with 1.4 ml methanol. To measure iron concentrations in the eluate, 50 µl of eluate was measured with an colorimetric iron detection kit (Sigma-Aldrich, Cat. No. MAK025-1KT) following the provided protocol. Iron concentration was determined by measuring absorbance at 595 nm on a plate reader.

#### Iron-limited *E. pacifica* pellets

were prepared by growing *E. pacifica* in 4% PG for 72 hours at 225 rpm and 30°C, then pelleted, aliquoted, and frozen into 10 mg pellets, and stored at −80°C. *E. pacifica* pellets were resuspended in 1ml of media to achieve a 10mg/ml liquid bacteria stock to add to cultures.

#### Culturing with different iron sources

To set up the iron utilization assay, cultures grown in 25% RA were passaged at a 1:60 dilution into 25% RA and grown for 72 hours at 18°C. This culture was then passaged at a 1:60 dilution into 25% RA and grown for 24 hours at 27°C. Finally, this culture was passaged at a 1:30 dilution into 4% PG (which had no iron added) and grown at 27°C for 24 hours. Cells from this final passage were twice washed by first centrifuging the culture at 2600xg for 4 minutes and then resuspending the pellet in 30 ml sterile ASW. After the final wash, the pellet was resuspended in 100 µl ASW and counted. These washed cells were used to seed cultures at 10^4^ cells/ml in 4% PG with the additions of 100 µM iron (ferric EDTA or ferric colloids) and 50 µg/ml of iron-limited *E. pacifica* pellets (see above). The resulting culture was grown at 27°C for 48 hours and harvested for each experiment.

### qPCR

**Protocol Links:** dx.doi.org/10.17504/protocols.io.ewov1nnzogr2/v1 dx.doi.org/10.17504/protocols.io.n2bvjnjongk5/v1

#### Primer design

Primers were designed against *cytb561a,* the experimental gene of interest and *cofilin,* a highly expressed eukaryotic specific housekeeping gene with a consistent expression between *S. rosetta* cell types, as shown by the RNAseq analysis. Primers for quantitative polymerase chain reaction (qPCR) were designed by selecting 19-20 bp oligos spanning exon-exon junctions from cDNA sequences, (downloaded from: https://protists.ensembl.org/Salpingoeca_rosetta_gca_000188695/Info/Index). Primer sequences as follows: *cytb561a* forward primer 5’-CATGGAAGAAGCGCATGGTG-3’, reverse primer 5’-GGGTCCGCAAGATCATTGAG-3’, *cofilin* forward primer 5’-CAAGCTCCCCACGGACAAG-3’, reverse primer 5’-GGGTCCATGCGAAGAAGAC-3’. In the case of *cytb561a,* primers were selected downstream of the premature termination sequence edited in by CRISPR/Cas9.

#### Primer validation

Target amplicons were amplified by polymerase chain reaction (PCR) from the cDNA of slow swimmer and thecate cultures (Fig. S4A) with the Luna Universal qPCR Master Mix (NEB, Cat, No. M3003). The amplicons were run on a 1% (w/v) agarose gel in Tris-Borate-EDTA (TBE) buffer to verify that the primer sets amplified only one amplicon. The amplification efficiency of primer sets was characterized across a serial dilution of ssDNA standards for each target (Fig. S4B-C). ssDNA standards were generated from PCR products amplified with a forward primer that was 5’ phosphorylated to promote Lambda exonuclease digestion of that strand (NEB, Cat. No. M0262S) and a reverse primer with phosphorothioate bonds between the first four 5’ nucleotides to block digestion. Those PCR products were then digested with Lambda exonuclease following NEB’s protocol. ssDNA concentrations were determined by Qubit ssDNA Assay Kit (Thermo Fisher, Cat. No. Q10212) by creating a standard curve with the provided standards and then reading fluorescence (Ex = 500 nm / Em=540 nm) on a plate reader (Molecular Devices, SpectraMax iD5). The ssDNA standards were serially diluted from 10^5^ to 10^0^ copies/µl in a solution with 10 ng/µl of *S. rosetta* RNA (Total RNA after removing mRNAs) to account for any matrix effects.

#### cDNA preparation

Samples from slow swimmer or thecate cultures were pelleted and lysed using the preferential lysis protocol, and total RNA was purified by RNeasy MinElute Cleanup Kit (Qiagen, Cat. No. 74204). cDNA was prepared by SuperScript IV Reverse Transcriptase (Thermo FIsher, Cat. No. 18090010) following the provided protocol using oligo d(T)_20_ for synthesis. However, the incubation temperature for cDNA synthesis was increased to 60°C to account for the high GC content of the *S. rosetta* genome.

#### qPCR

3 µl of cDNA samples or ssDNA standards were added to a 20µl reaction with Luna Universal qPCR Master Mix (NEB, Cat, No. M3003) according to the provided protocol. The samples were amplified in a single run on the QuantStudio 3 Real-Time PCR System (Thermo Fisher, Cat. No. A28567).

### Genome editing

**Protocol Link:** dx.doi.org/10.17504/protocols.io.j8nlk86o5l5r/v1

#### Culturing cells for transfection

Mutant strains were generated with a modified protocol from a previously published method^46^. Cultures of *S. rosetta* slow swimmers were maintained in 15% RA/2% PYG (called 15/2, Table S1) at 22°C, and 48 hours prior to transfection, cultures were seeded at 10^4^ cells/ml in 80 ml of 15/2 Media in 300 cm^2^ vented tissue culture flasks (VWR, Cat. No. 10062-884) and grown at 22°C.

#### Cas9 RNP preparation

On the day of transfection, Cas9 bound to a guide RNA (Cas9 RNP) was prepared by combining 2 µl of 100 µM guide RNA (gRNA) with 2 µl of 20 µM EnGen SpyCas9 NLS (NEB, Cat. No. M0646T), and incubated at room temperature for 2-4 hours. (Note: Our gRNA was ordered as synthetic RNA oligonucleotides from Integrated DNA Technologies that came as an Alt-R CRISPR-Cas9 crRNA and an Alt-R CRISPR-Cas9 tracrRNA. These two oligonucleotides were annealed together to form gRNA). Guide sequences can be found in table S4.

#### Transfection

During the Cas9 RNP incubation, cells were harvested for transfection by centrifuging the cultured cells at 2400xg for 3 minutes at 4°C. The supernatant was discarded, and cells were resuspended in 50 ml of Cell Wash Buffer (420 mM NaCl; 50 mM MgCl_2_; 30 mM Na_2_SO_4_; 10 mM KCl; titrated to pH 8.0 with ∼2.4 mM NaHCO_3_). The cells were centrifuged again at 2400xg and 4°C for 3 minutes. The supernatant was removed, and the pellet was resuspended in 100 µl of Cell Wash Buffer. Cells were counted and diluted to 5×10^7^ cells/ml, split into 100 µl aliquots, and centrifuged for 2 min at 800xg and room temperature. The supernatant was removed and replaced with 100 µl priming buffer (40 mM HEPES-KOH, pH 7.5; 34 mM Lithium Citrate; 15% (w/v) PEG 8000; 50 mM Cysteine; 1.5 µM Papain). Cells were primed for 45 minutes at room temperature. Afterwards, the priming was quenched with 10 µl of 50 mg/ml bovine serum albumin. The primed cells were centrifuged at 1200xg for 4 min at room temperature. After removing the supernatant, the cells were resuspended in 25 µl ice cold SF buffer (Lonza Cat. No. V4SC-2096). Nucelofection reactions were prepared by combining 16 µl ice cold SF buffer, 4 µl Cas9 RNP, 2 µl of repair oligo, and 2 µl of the resuspended and primed cells. This reaction mix was loaded into a 96-well nucleofection plate (Lonza Cat. No. AAF-1003S, AAF-1003B), and pulsed with CU 154 (Fig. S4). 100 µl of ice cold recovery buffer (10 mM HEPES-KOH, pH 7.5; 900 mM Sorbitol; and 8% (w/v) PEG 8000) was added immediately to each pulsed well and incubated for 5 minutes at room temperature. Afterwards the entire content from each nucleofection well was transferred into 2 ml of 15/2 Media with the addition of 50 µg/ml *E. pacifica* pellet.

### Strain isolation and Screening

**Protocol Link:** dx.doi.org/10.17504/protocols.io.14egn6n2pl5d/v1

We adapted the Cas12a DETECTR genotyping assay^101^ to screen for cells with desired mutations from Cas9 genome editing. Following overnight recovery post-nucleofection, cells were counted and diluted to 45 cells/ml in 10% RA (Table S1) with 50 µg/ml *E. pacifica* pellet and aliquoted into 96-well plates. Plates were then grown at 27°C for 72 hours to propagate cells. Afterwards 12 µl from each well was added into 36 µl DNAzol Direct (20 mM potassium hydroxide; 60% (w/v) PEG 200, pH 13.3-13.7– Note: It is important to test a range of pH values to establish the optimal pH for your own use)^46^ that was pre-aliquoted into 96-well PCR plates, which we called the lysed sample. The lysed samples were then incubated at 80°C for 10 minutes. The target DNA was amplified (primer sequences in Table S4) in PCR reactions prepared with GoTaq Clear Master Mix (Promega, Cat. No. M7123) using 2 µl of the lysed samples per 25 µl PCR reaction. Cycling conditions followed Promega guidelines and melt temperatures for oligos were calculated using (https://www.promega.com/resources/tools/biomath/tm-calculator/). During the PCR, a Cas12a mastermix was assembled in two parts: First, Cas12a RNP: (13 µl water; 2 µl r2.1 buffer [NEB, Cat. No. B6002S]; 2.5 µl 100 µM gRNA [ordered as a synthesized oligo]; 2 µl 100 µM LbaCas12a [NEB, Cat. No. M0653T]) was incubated for 5 minutes at room temperature.

Second, the Cas12a RNP was combined with the rest of the components (for one 96-well plate: 486 µl water; 60 µl r2.1 buffer; 18 µl Cas12a RNP; 36 µl 5 µM ssDNA probe [IDT, Cat. No. 11-04-02-04]) and then incubated for 5 minutes at room temperature. 5 µl of the mastermix was added to each 25 µl PCR and incubated at 37°C for 1 hour. The fluorescent signal was then measured in the VIC channel on a QuantStudio 3 Real-Time PCR System. Wells with high signal (≥10-fold above background) were recovered in 25% RA overnight at 27°C, counted, diluted to 3 cells/ml in 10% RA with 50 µg/ml *E. pacifica* pellet, and plated in a 96-well plate with 100 µl/well, which corresponds to 0.3 cells/well. Plates were grown at 27°C for 72-96 hours. Finally, these clonal isolates were screened by digesting amplicons with a restriction enzyme (RE). 12 µl per isolate was added to 36 µl DNAzol, incubated at 80°C to make a lysed sample. 2 µl of the lysed samples were added to 25 µl PCR reactions with GoTaq Green mastermix (Promega, Cat. No. M7123). After amplification, 1 µl of RE was added to each 25 µl PCR reaction, and incubated following manufacturers’ protocols. RE digestions were run on 1% agarose gels with wild-type or undigested controls. For positive RE digestion hits, PCR products were Sanger sequenced to confirm the sequence of the edited site (Table S5).

### Growth curves

#### S. rosetta

We monitored cell density at times points over 48 hours to assess the population dynamics of wild type and *cytb561^PTS^* thecates in iron utilization assay conditions. Cells were plated in 24-well plates with 100 µM ferric EDTA or ferric colloids sources in 4% PG or in nutrient replete media (25% RA) as a control. At each time point, the entire contents of one well was harvested by scraping the bottom and then transferring the liquid to a microfuge tube. Cells were fixed with 5 µl of 37.5% paraformaldehyde (PFA) and then counted on a Reichert Bright-Line Metallized Hemacytometer (Hausser Scientific, Cat. No. 1483). Growth curves were fit to a logistic growth equation that explicitly models the lag time with a heaviside step function^28^.

#### E. pacifica

growth curves were set up in 4% PG media inoculated with 50 µg/ml iron-limited *E. pacifica* pellet and a ferric EDTA titration in clear bottom 96-well plates, and grown for 48 hours at 27°C. Growth was assessed by monitoring the optical density at 600 nm on a plate reader with orbital shaking before every reading. Ferric colloids were not included in the growth curves as iron particulates would shift optical density readings. To assess *E. pacifica* growth with ferric colloids, 4% PG media was inoculated with 50 µg/ml iron-limited *E. pacifica* pellet and 100 µM ferric EDTA or ferric colloids in 6-well plates and grown for 48 hours at 27°C. Wells were scraped and 1 ml aliquots were centrifuged at 500xg for 2 minutes to settle iron particulates. The top 500 µl of supernatant was gathered for measuring the optical density at 600 nm.

### Immunofluorescent Staining

**Protocol Link:** dx.doi.org/10.17504/protocols.io.5jyl85k38l2w/v1

Cultures of wild-type and ALFA-tagged strains were maintained in 25% RA at 18°C for thecates and 22°C for slow swimmers, as higher temperatures helped to maintain slow swimmers. 24 hours prior to imaging, cultures were seeded at 10^4^ cells/ml in 6 ml of 25% RA in 25cm^2^ vented tissue culture flasks (VWR Cat. No. 10062-874) and grown at 27°C. On the day of imaging, prior to preparing cells, 50 µl of 10 mg/ml Poly D-lysine hydrobromide (MP Biomedicals, Cat. No. 102694) was added to each well of a glass bottom 96-well plate (Thermo Scientific, Cat. No. 165305) and incubated for ≥15 min at room temperature. To prepare cells, the supernatant of thecate cultures was decanted, and replaced with 6 ml filtered ASW, then scraped with a cell scraper, and centrifuged at 500xg for 5 minutes at room temperature. For swimmer cultures, the cells were centrifuged 500xg for 5 minutes at room temperature. The supernatant removed leaving 0.5 ml remaining, and resuspended in 5 ml filtered ASW and centrifuged 500xg for 5 min at room temperature. Both thecate and swimmer cultures follow the same protocol henceforth. The supernatant was removed, leaving 0.5 ml, and the tubes gently swirled to resuspend cells.

The poly D-lysine coated wells were washed 3 times with 50 µl ASW, and then 50-100 µl of cells were added to each well using wide-bore pipette tips. Cells settled on the well surface for 15 minutes. All subsequent steps use gel loading tips. After settlement, the supernatant was removed leaving 25 µl behind to avoid drying and damaging delicate cell structures. 70 µl of Fix buffer (10 mM MES-KOH, pH 6.1; 138 mM KCl; 3 mM MgCl_2_; 2 mM EGTA; 3% (v/v) PFA; 15% (w/v) sucrose) was slowly added to the top of each well and incubated for 5 min at room temperature. 70 µl of supernatant was removed from the opposite side from which solution was added. This will be referred to as a “wash”. Next, wells were washed with Fix/tween buffer (Fix buffer with the addition of 0.07% (v/v) Tween-20) and incubated for 5 min at room temperature. Wells were then washed with a quench buffer (10 mM MES-KOH, pH 6.1; 138 mM KCl; 3 mM MgCl_2_; 2 mM EGTA; 15% (w/v) sucrose; 0.3M glycine) and washed immediately with Permeabilization-MeOH buffer (8% (v/v) MeOH; 1% (v/v) Tween-20; in LICOR Intercept Blocking Buffer [LI-COR Biosciences Cat. No. 927-70001]) and incubated for 15 minutes. (Note: LICOR Intercept Blocking buffer increased detection of AFLA epitopes during testing.) After, wells were washed with Permeabilization buffer (1% (v/v) Tween-20; LICOR Intercept Blocking Buffer) and then washed twice in antibody mix (1:200 FluoTag-X2 anti-ALFA Alexa647 [Nanotag, Cat No. N1502-AF647-L]; 1:200 phalloidin Alexa546 [Invitrogen, Cat. No. A22283]; 1:500 alpha Tubulin Monoclonal Antibody, DM1A [Invitrogen, Cat. No. 62204]; 1:500 alpaca anti-mouse IgG1, recombinant VHH, Alexa488 [Chromotek, Cat. No. sms1AF488-1]) diluted in the Permeabilization buffer and incubated for 1 hour protected from light. Afterwards, wells were washed twice with PEM (100 mM PIPES-KOH, pH 6.9; 1 mM EGTA; 1 mM MgSO_4_) and immediately imaged.

### Microscopy

#### Time-lapse of ferric colloid ingestion

100 µl of 1.46 µM fluorescent ferric colloids were added to 2 ml of wild-type thecate cultures, previously grown in glass bottomed dishes (World Precision Instruments, Cat. No. FD35-100) for 24 hours at 22°C. Samples were imaged by widefield microscopy. The microscope was a Nikon Eclipse Ti2-E inverted microscope outfitted with a D-LEDI light source, Chroma 89401 Quad Filter Cube, and 60x CFI Plan Apo VC NA1.2 water immersion objective. Time-lapse images were acquired every 4 sec over 2 hours with 50 msec exposure time for brightfield and fluorescent (TRITC channel) images. In three independent experiments, a total of 4 feeding events were found in a field of 200 cells, although this was not an exhaustive search for ferric colloid intake frequency. One cell that was oriented on its side to best visualize feeding was selected for further processing. Images were processed in FIJI by cropping individual cells in 100×100 pixel bounds, and image contrast was manually adjusted. Ferric colloid particles were automatically tracked, first by setting a threshold to produce a binarized image (image > adjust > manual threshold) and then by finding particles (analyze > analyze particles > size = 10-300 pixels, circularity = 0.2-1.0).

#### Immunofluorescence Imaging

Samples were imaged by widefield microscopy. The microscope was a Nikon Eclipse Ti2-E inverted microscope outfitted with a D-LEDI light source, Chroma 89401 Quad Filter Cube, and 60x CFI Plan Apo VC NA1.2 water immersion objective. Single focal plane and Z-stacks of samples were imaged with two-fold binning (120 µm/pixel) on a Nikon Digital Sight 50 M Camera with 100-400 msec exposure times for each channel and variable illumination intensities to extend the dynamic range of signal without photobleaching samples. Single focal-plane images were processed within the Nikon imaging software. Images were background subtracted (120 pixel, rolling ball radius), and deconvolved using the Landweber deconvolution algorithim with 12-16 iterations (autostopping engaged), and automatic background subtraction. Deconvolved images were cropped in 150×150 pixel bounds in FIJI and merged images were combined using maximum intensity composite. Supplementary images were processed in FIJI by cropping individual cells in 150×200 pixel bounds, projecting the maximal intensity, subtracting the background (75 pixel, sliding paraboloid radius), enhancing the contrast (saturation 0.02% of the pixels), and applying a minimum filter of 0.5 pixels. Merged images were combined using maximum intensity composite.

### Western blots

**Protocol Link:** dx.doi.org/10.17504/protocols.io.3byl4987zgo5/v1

Cultures were grown in 25% RA or 4% PG (without supplemented iron) for 24 hours at 27°C to test whether external iron conditions (replete or deplete) resulted in any post-translational regulation of protein expression. Cultures were scraped and then lysed according to the preferential lysis protocol, with the following modifications: TurboDNase and SUPERasein were replaced with 15 µl of Pierce universal nuclease (Thermo Scientific Cat. No. 88702) per 1 ml buffer, and heparin and cycloheximide were removed. After clearing the lysate, Tween-20 was added to a final concentration of 1% (v/v), and the sample was incubated for 10 minutes at room temperature to aid in liberating membrane-bound proteins. Protein concentration was determined by Bradford assay (Thermo Fisher Scientific Cat. No. 23236) and samples were normalized to the lowest concentration. To prepare for SDS-PAGE, approximately 20 µg or 40 µg of total protein for thecates or slow swimmers, respectively, was denatured with 4x Laemmli SDS loading buffer and incubated for 10 minutes at room temperature (Note: 80°C caused membrane proteins to crash out, so room temperature incubation was essential). Samples were loaded into a Novex wedgewell 10% Tris-Glycine Mini Gel (Thermo Fisher Scientific Cat. No. XP00105BOX) along with a 1:10 diluted PageRuler Plus Prestained Protein Ladder (Thermo Fisher Scientific Cat. No. 26619). Gels were run at 225 V for 30 minutes, and afterwards the gel was transferred to a 0.45 µM nitrocellulose membrane (Bio-rad Cat. No. 1620115) at 50 V for 30 minutes, then 100 V for 60 minutes in transfer buffer (25 mM tris base; 192mM glycine; 10% methanol) at 4°C. Afterwards the membrane was stained with total protein stain (LI-COR Biosciences Cat. No. 926-11010) and then imaged in the 700 nm channel with a 30 second exposure on a LI-COR Odysey Fc. Afterwards, the membrane was blocked in LICOR Intercept Blocking Buffer (LI-COR Biosciences Cat. No. 927-70001) for 1 hour at 37°C and then overnight at 4°C. The following day the membrane was immunostained with FluoTag-X2 anti-ALFA LI-COR IRDye 800CW (NanoTag Biotechnologies Cat. No. N1502-Li800-L) diluted 1:10,000 in LI-COR PBS block plus 0.2% Tween-20 for 2 hours at room temperature. (Note: Both the LI-COR PBS block and anti-ALFA IRDye 800CW were essential for visualization, other blocks and fluorophores lead to significantly lower signal.) The membrane was washed 3x in PBST (PBS with 0.2% Tween-20) and then imaged in the 800 nm channel with a 10 minute exposure. This western protocol was adapted from a previous study^95^.

### Protein database searches and tree construction

We compiled a list of iron acquisition proteins (Table S6) from these model organisms: *Homo sapiens*^44^, *Saccharomyces cerevisiae*^47^, *Chlamydomonas reinhardtii*^43^, *Arabadopsis thaliana*^102^, and *Phaeodactylum tricornutum*^41^. For each protein, we retrieved the corresponding Pfam hidden markov model (HMM) from Interpro (https://www.ebi.ac.uk/interpro/)^103^. Using HMMER v3.4 (hmmer.org^104^), we searched proteomes from EukProt v3^105^, focusing on proteomes from “The Comparative Set” (TCS), which consisted of 196 species of phylogenetic relevance with high BUSCO completion. We supplemented the TCS with the transcriptomes of 18 choanoflagellate species^54^ and other Holozoa and Fungi. To count the number of unique proteins that conformed to the HMM profile from ‘hmmsearch’, we aligned hits with ‘hmmalign’ and retained only one sequence from hits with >99% identity (Note: this was important for removing multiple isoforms and reference sequences from highly annotated genomes). We additionally filtered hits based on length with ‘esl-alimanip’ to retain protein sequences that only possessed the HMM profile, which eliminated multidomain proteins that only had the HMM profile as a one of its domains (Table S3 for length filter thresholds). (Note: The number of hits prior to filtering can be found in Table S3) For HMM profiles that yielded a high number of protein hits that were not well-filtered (e.g. Transferrin Receptor), we aligned sequences with MAFFT-DASH^61^, removed sites that were uninformative or contained 90% gaps with ClipKIT^106^ (kpic-gappy 0.9), and generated phylogenetic trees with IQ-TREE^107^ using ModelFinder^108^ to choose the appropriate substitution model for each set of protein alignments. This procedure was also used to determine the phylogenetic relationships of Cytb561 paralogs (Fig. 4A), and the chosen substitution model was LG+F+R7. Trees were visualized with iTOL^109^. Multiple sequence alignments were visually inspected and curated for supplemental figures using Jalview v2^110^. Cytb561 structure models were predicted with AlphaFold v3^65^ and visualized using UCSF ChimeraX^64^.

### Gene distribution maps and correlations with upwellings

#### *cytb561* paralog search

The abundance and location of *S. rosetta* cytb561 paralogs was searched for using the Ocean Gene Atlas^8,9^. Protein sequences of each paralog from *S. rosetta* were searched for in the MATOUv1+G (for metagenomes) and MATOUv1+T (for metatranscriptomes) databases, using tblastn with the following search parameters: expected threshold E-values < 1^-10^, and abundance as percent of total reads. The Ocean Gene Atlas automatically assigns protein sequences to phyla and classifies them on expected organism size, such that the results could be filtered first by size (0.8-5 µm) and then by phyla to identify only choanoflagellate specific hits. Therefore to identify choanoflagellate specific hits and the location they were gathered from, three tables were gathered from each search: the distribution of E-values (E-values per hit and their assigned phyla), abundance matrix for the 0.8-5 µm fraction (the abundance as a fraction of total reads and the sample site ID for each hit), and environmental parameters for the 0.8-5 µm fraction (all measured and inferred environmental measurements for each hits sample location). From these three tables, choanoflagellate assigned hits, abundances, and sample locations were compiled.

#### Upwelling velocities

To identify average vertical velocities at the sample sites from the Tara Oceans Expedition^111^, from which the Ocean Gene Atlas acquires its data, upward seawater velocities were gathered from the Copernicus Marine Services’ Global Ocean Physics Analysis and Forecast model’s (https://doi.org/10.48670/moi-00016) dataset (cmems_mod_glo_phy-wcur_anfc_0.083deg_P1M-m). The coordinates for each sample site were compiled^111^ and vertical velocities were gathered from the dates 10/31/2020 - 5/20/2024. Then the search was done using a modified script offered by the Copernicus Marine Services (https://help.marine.copernicus.eu/en/articles/7970637-how-to-download-data-for-multiple-points-from-a-csv). The vertical velocity at each sample site was then averaged across the 3.5 year range to account for temporal variation (Fig. S8B).

#### Correlations and maps

Choanoflagellate paralog abundances were tabulated with their sample site ID, sample coordinates, and average upwelling velocity (from above). First, the abundances and average vertical velocities were correlated by the Pearson function in Microsoft Excel. To calculate the associated *P* value for each Pearson correlation, a T statistic was calculated using the formula: *t = (r•[n-2]^½^)•(1–r^2^)^-½^*, where *r* is the calculated Pearson correlation, *n* is the sample number, *t* is the T statistic. This statistic was then input into the t distribution function (TDIST) in Excel, along with the degrees of freedom (*n*-2), to calculate the *P* values. Next, the abundances and average vertical velocities were plotted in a scatter plot to manually inspect the data for outliers. If a datum was suspected of being an outlier, a Dixon’s Q test was conducted for that datum with the formula: *Q = |gap|/range,* where gap = the difference between the suspected outlier and the next nearest datum, and range = the range of the data set. The value is considered an outlier and removed if Q > Q_table_, where Q_table_ is a reference value determined by the sample size and confidence level. Of all tested outliers, only one datum was removed, belonging to *cytb561a* transcript abundance at TARA site 52 (−16.96°, 53.99°), with a Q = 0.8718 and where Q_table_ = 0.3323 for 99% confidence level for a sample size of 33. Without this outlier removed the statistics for *cytb561a*’s correlation would be r = 0.09 and *P* = 0.58. Maps were plotted with the compiled table of paralog abundances and coordinates, using matplotlib with modified scripts generated by ChatGPT (https://chat.openai.com).

## Data Availability

RNA sequencing data is available in the public repository NCBI Gene Expression Omnibus (GEO) database under requisition ID: GSE267344

## Author Contributions

FL and DSB conceived of the project, designed experiments, conducted research, performed data analysis, and wrote the manuscript. JMEE and VD generated strains and developed the genotyping pipeline. MCC and SE performed an early analysis of RNA-seq data.

## Acknowledgements

We thank Nicole King for supporting an early phase in this project and providing feedback. Alain Garcia de las Bayonas, Flora Rutaganira, Ben Larson, members of the Booth lab, and the students at the Marine Biological Laboratories (MBL) 2023 Physiology Course provided essential experimental advice. We also appreciate helpful discussions from Elizabeth Blackburn, Jeremy Reiter, Wallace Marshall, Sandy Johnson, Carol Gross, and Dyche Mullins. FL was supported by a National Science Foundation Graduate Research Fellowship, a UCSF Discovery Fellowship, and the UCSF Tetrad Graduate Program Training Grant T32GM139786. This work was supported by awards to DSB from the Chan Zuckerberg Biohub, the UCSF Sandler Program for Breakthrough Biomedical Research, a David and Lucile Packard Foundation Fellowship in Science and Engineering, and a Maximizing Investigators’ Research Award for Early Stage Investigators from the National Institute of General Medical Sciences R35GM147404. Sequencing was additionally supported by an instrumentation grant from the National Institutes of Health S10OD018174 to the QB3 Vincent J. Coates Genomics Sequencing Laboratory at University of California, Berkeley.

