## Supplemental Figures & Table Legends for "Cell differentiation controls iron assimilation in a choanoflagellate"

**Title:**

**Affiliations:**

[3]Current Address: Department of Molecular Biosciences
University of Texas, Austin
Austin, TX 78712

**Supplemental Figures:**

**Figure S1.** A preferential lysis buffer enriches RNA from *S. rosetta* to improve transcriptomes.
**(A)** A schematic showing the separation of *S. rosetta* RNA from *E. pacifica* RNA.
**(B)** The preferential lysis procedure enriches RNA from *S. rosetta*. After performing preferential lysis, RNA was purified from the supernatant using Trizol LS reagent. The RNA from the pellet of preferential lysis, an unlysed sample of *S. rosetta* feeding on *E. pacifica*, or a sample of *E. pacifica* was purified with Trizol LS. The purified RNA samples were combined with an equal volume of formamide and incubated at 95°C for 5 minutes. After cooling to room temperature, the samples were run on a 2% (w/v) agarose gel in TBE buffer at 100 V. Afterwards, the gel was stained with SYBR Gold to detect RNA. In the gel, the supernatant (S) from whole lysates (W) mostly contains *S. rosetta* rRNA (28S and 18S); the pellet (P) contains E. pacifica rRNA (23S and 16S).
**(C)** Total RNA prepared for RNA-seq primarily contains rRNA from *S. rosetta.* Bioanalyzer traces with predicted prokaryotic and eukaryotic rRNA subunit sizes indicated. Replicate RNA samples used for sequencing demonstrate little to no carryover from bacterial contamination, as shown by the low amount of 16S and 23S rRNA.

**Figure S2:** GO molecular functions and biological processes enriched in thecates. Functional modules enriched in thecates mediate signal transduction, nutrient acquisition, and gene regulation. Genes with transcript abundances that reliably ($q < 0.01$) changed more than two-fold in thecates were analyzed for the enrichment of Gene Ontology (GO) categories, focusing on GO molecular functions **(A)** and biological processes **(B)**. Genes that were upregulated in thecates and associated with an enriched GO molecular function category were counted (x-axis). Each bar on the graph is colored by the *P*-value, which reflects the probability that a given category was represented by a random selection of genes from the *S. rosetta* genome.

**Figure S3:** GO molecular functions and biological processes enriched in swimmers. Functional modules enriched in swimmers support RNA processing, mitosis, and metabolism. Genes with transcript abundances that reliably ($q < 0.01$) changed more than two-fold in swimmers were analyzed for the enrichment of Gene Ontology (GO) categories, focusing on GO molecular functions **(A)** and biological processes **(B)**. Genes that were upregulated in swimmers and associated with an enriched GO molecular function category were counted (x-axis). Each bar on the graph is colored by the *P*-value, which reflects the probability that a given category was represented by a random selection of genes from the *S. rosetta* genome.

**Figure S4:** Validated qPCR primers confirm the disruption of *cytb561a* expression with the *cytb561a*$^{PTS}$ allele.

**(A)** qPCR primers for *cytb561a* and *cofilin* produce a single amplicon*.* Gel image of PCR products amplified from qPCR primers designed for *cytb561a* and *cofilin*. Note the diminished band from swimmer cDNA likely resulted from lower mRNA copies from slow swimmer cultures, as shown in Fig. 2C. Shadow at 75 bp is a shadow of the loading dye.

**(B)** Standard curve for qPCR primers for *cytb561a*. To generate a standard curve, a serial dilution of ssDNA standards was performed in triplicate. The concentration of the ssDNA standard was determined by Qubit. Cycle thresholds (Ct) were converted to transcript copy number by fitting a linear equation to the standard curve. This curve was used to determine the abundance of transcripts from three independent experiments.

**(C)** Standard curve for qPCR primers for *cofilin*. The graph is laid out as for panel B.

**(D)** *Cytb561a*$^{PTS}$ exhibits lowered *cytb561a* expression compared to wild-type thecates. We compared the normalized expression of *cytb561a* from the strain bearing the *cytb561a*$^{PTS}$ allele to wild-type strains. The wild-type strains were slow swimmers cultured with ferric EDTA or thecates cultured with ferric colloids, for we anticipated that these two conditions would result in the greatest difference in *cytb561a* expression.

**Figure S5:** *S. rosetta* cell types, but not their feeder bacteria, grow differently with variable iron sources.

**(A)** Thecate cell types exhibit differential growth with low concentrations of ferric colloids compared to slow swimming cell types. *S. rosetta* cell density of slow swimmer and thecate cultures when grown with titrations of ferric EDTA or ferric colloids.

**(B)** A variety of ferric chelates differentially impact *S. rosetta* growth. Slow swimmers and thecates exhibit different growth characteristics, so to better compare their growth with iron bound to different chelators, we used a ratio of the final cell density of cultures grown with iron-chelator complexes to the final cell density of cultures grown without any supplemental iron. With this metric, a ratio greater than one indicates that the cell type displays increased growth with the iron-chelator complex; whereas, a ratio less than one indicates the converse. All conditions were tested at 100 μM $Fe^{3+}$, and pyoverdines and deferroxamine are bacterial siderophores.

**(C)** *E. pacifica* feeder bacteria growth is easily rescued by low concentrations of iron. Optical density at 600 nm (OD600) of *E. pacifica* measured approximately every 100 minutes for 48 hours. Note standard deviations overlap for all conditions except for a no iron control.

**(D)** *E. pacifica* grows similarly with ferric EDTA or ferric colloids. Because ferric colloids would influence OD600 measurements in a standard growth curve experiment, *E. pacifica* cultures were grown for 48 hours with 100 µM ferric EDTA or ferric colloids and then lightly centrifuged at 500xg for 2 minutes at room temperature to settle iron particulates. Afterwards, the supernatant OD600 was measured.

**(E)** Ferric colloids release a small amount of labile iron. When media is prepared with 100 µM ferric colloids, only 3.43 ± 1.01 µM iron is liberated without biological activity. The amount of iron in 4% PG media alone was 0.27 ± 0.036 µM iron.

**Figure S6:** A screen for optimal nucleofection pulses to improve cargo delivery into *S. rosetta*.

**(A)** Nanoluc luminescence values from *S. rosetta* cultures transfected with different pulse codes. Because of changes to the growth media for *S. rosetta*, we conducted a new screen for an optimal nucleofection pulse. Cells were transfected with a plasmid that drives nanoluc expression with a promoter cloned from elongation factor L. Pulse codes correspond to Lonza 4D-Nucleofector codes. The screen was performed in four independent trials. Dot indicates prior pulse code used, asterisk denotes optimized pulse code.

**(B)** Re-ordered chart of luciferase activity from the pulse screen reveals the landscape to optimize nucleofection pulses. After the first two trials for screening pulses, we hypothesized that the letter and number codes for the nucleofection system are orthogonal parameters, so we selected pulses for the third and fourth trials to more broadly sample the optimization landscape. The maximum luminescence value for each pulse from one of the four trials was used to visualize transfection efficiency as a function of the letter codes (x-axis) or pulse numbers (y-axis). Dot indicates prior pulse code used, asterisk denotes optimized pulse code.

**Figure S7:** Genotypes of engineered strains.

For each strain that was engineered, the targeted locus (unique identifiers and allele names shown on the left) was amplified by PCR and then sequenced with the Sanger method. The sequencing traces for locus are displayed with the predicted and confirmed sequences. Two strains ($heph^{alfa}$ and $cytb561c^{alfa}$) were heterozygous with a wild type and edited allele, as seen in superimposed peaks at each nucleotide position. The sequences of heterozygotes were further analyzed by DECODR[112] to confirm the presence of both sequences.

**Figure S8:** Thecates grow slight faster and to a larger population size with ferric colloids

**(A-C)** Growth curves of wild type and *cytb561a*$^{PTS}$ thecates show wild type thecates grow to a larger population size when grown with ferric colloids. Growth curves of cultures grown in (A) nutrient replete conditions (25% RA), (B) 4% PG with ferric EDTA, and (C) 4% PG with ferric colloids. Cell densities were counted every 6 hours for 48 hours.

**(D-E)** Comparisons of growth curve parameters highlight the increase in carrying capacity as the main factor for higher wild type thecate cell type growth with ferric colloids. (D) Carrying capacity, and (E) doubling time of the growth curves shown in figures (B) and (C).

**Figure S9:** Thecates phagocytose ferric colloids.

**(A)** Fluorescently labeled ferric colloids were embedded with fluorescent dextrans. During the precipitation of $FeCl_3$ to make ferric colloids, fluorescently labeled dextrans were added. The resulting colloids were incubated with dextranase, washed, and then resuspended in ASW before adding to thecate cultures for time-course microscopy.

**(B)** Embedded fluorescent dextrans highlight particles of ferric colloids. The images show colloids of similar size and density, with only the co-precipitated ferric colloids and dextran exhibiting fluorescence.

**(C)** Corroborating examples of ferric colloid ingestion. Time courses of thecates feeding on ferric colloid particles (magenta) are displayed in the fluorescence channel (below) and in a merge with brightfield and fluorescence channel. 0 seconds denotes the initial point of contact between the cell and the tracked colloid particle.

**Figure S10:** Simplified iron acquisition pathways in model eukaryotes.

**(A)** Mammals. Dietary iron is absorbed by the epithelium of the gut. Ferric cations are reduced and imported at the apical end of epithelial cells. Ferrous cations are exported and oxidized at the basal end into the bloodstream. Ferric cations are bound to transferrin in the bloodstream and distributed to other cells in the body by receptor mediated endocytosis when transferrin binds to the transferrin receptor.

**(B)** *Saccharomyces cerevisiae.* In the main iron import pathway, ferric cations are reduced, then oxidized, and finally imported in its ferric form for increased substrate specificity. In a minor secondary pathway ferrous cations can be imported in its reduced form by a low affinity importer. Iron is stored intracellularly in vacuoles, via another iron specific transporter.

**(C)** *Phaeodactylum tricornutum.* Ferric cations can be reduced directly and imported, but they can also be directly bound by phytotransferrin and endocytosed.

**(D)** *Chlamydomonas reinhardtii.* Ferric cations are first reduced, then oxidized, and finally imported in its ferric form for increased substrate specificity. Iron can also be directly bound by phytotransferrin, which is thought to be released into the periplasm.

**Figure S11:** Phylogenetic trees of iron acquisition proteins

Unrooted maximum likelihood phylogenies of the iron acquisition proteins shown in Fig. 3A. Alignment, trimming, and tree building details are listed in table S3.

**Figure S12:** Animal transferrin receptors evolved from an ancestral M28 peptidase.

**(A)** Animal transferrin receptors emerged from ancestral animal M28 peptidase. Maximum likelihood phylogeny built from a search using a new HMM model of the domain architecture shown in (B). Animal transferrin receptors are highlighted by a gray outline, which was determined by the loss of peptidase enzymatic residues shown in (C) and (D).

**(B)** The transferrin receptor protein domain architecture. Domain labels are for the protease associated (PA) domain, M28 peptidase (M28 Pept.) domain,  and the transferrin receptor specific dimerization (Dimer) domain. Pfam accession IDs are listed below the associated domain.

**(C)** Transferrin - transferrin receptor complex with highlighted catalytic residues. The structure of transferrin - transferrin receptor complex as determined by cryo-electron microscopy (PDB 1SUV). The catalytic residues of the transferrin receptor are highlighted and colored.

**(D)** The loss of key catalytic residues marks the evolutionary transition from peptidase to transferrin. Alignments of transferrin receptors across animals show a remarkable loss of key M28 peptidase catalytic residues. This loss likely demarks the transition from peptidase to transferrin receptor, yet the binding of transferrin outside the catalytic site may have allow some related proteins to have dual functions as peptidases and transferrin receptors.

**Figure S13:** Thecates detectably express Cytb561a, Fpn, and Heph in western blots.

**(A)** Western blot of thecates cultures of ALFA-tagged strains, grown in nutrient rich media (25% RA) and iron depleted media (4% PG). Strains are named after the ALFA-tagged gene.

**(B)** Western blot of swimmer cultures of alfa-tagged strains, grown in nutrient rich media (25% RA) and iron depleted media (4% PG). Arrow indicates faint Fpn band visible only in the 25% RA condition. The expression of Fpn in media with high iron concentrations and the lack of expression in low iron media is consistent with Fpn regulating homeostatic iron levels through post-transcriptional regulation.

**Figure S14:** Thecates stably express Cytb561a independent of iron availability. Western blot of *cytb561a*$^{ALFA}$ thecates and swimmers grown in nutrient rich media (25% RA) and iron depleted media (4% PG) show minimal change is response to iron availability, indicating little to no post-transcriptional regulation of Cytb561a.

**Figure S15:** Additional immunofluorescence of thecates and swimmers

**(A)** The localization of Cytb561a, Fpn, and Heph in thecates, visualized through immunofluorescence of ALFA tagged proteins. Images are not deconvolved, and only processed in FIJI according to the methods.

**(B)** The localization of Fpn and Heph in swimmers, visualized through immunofluorescence of ALFA tagged proteins. Note, Cytb561a is not detectable in swimmers, matching the protein levels seen in western blots (Fig. S13) and by transcript levels shown by RNAseq and qPCR (Fig. 2B-C). Images are not deconvolved, and only processed in FIJI according to the methods.

**Figure S16:** Cytochrome b561 paralogs have retained ascorbate binding residues.

**(A-B)** The ascorbate binding pocket of human cytochrome b561 protein DCYTB. The crystal structure (PDB 5ZLG) of human duodenal cytochrome b561 (DCYTB) features a positively charged binding pocket on the cytosolic face of each polypeptide of the homodimeric complex **(A)**. Each pocket binds one ascorbate molecule (orange) that is coordinated by specific electrostatic and hydrophobic contacts **(B**, cyan residues**)**

**(C)** Cytb561 homologs have retained ascorbate binding residues. An alignment of cytochrome B561 protein sequences shows conservation in positions that correspond to ascorbate-binding residues in human DCYTB.

**Figure S17:** Cytochrome b561 dimerization is a feature of Group A proteins.

**(A-B)** The crystal structure of human DCYTB (PDB 5ZLG) forms a homodimeric complex **(A)**. Residues that mediate homodimerization **(B, pink residues)**. Heme groups are depicted in black with coordinated iron atoms in red.

**(C)** Cytochrome b561 homologs from Group A, including human DCYTB and *S. rosetta* Cytb561a, have key residues for dimerization, while groups B and C do not. An alignment of cytochrome b561 protein sequences reveals that Group A homologs conserve similar residues at positions that correspond to the dimerization residues in DCYTB. Sequences in Group B do not align at ~7 positions corresponding to the dimeric interface, and Group C sequences do not align at ~10 positions.

**Figure S18:** Cytb561a is predicted to dimerize similarly to human DCYTB.

**(A)** Expected position errors of dimerization predictions indicate that Cytb561a robustly forms a dimeric complex. Dimeric structures of each Cytb561 paralog bound to heme cofactors were predicted with AlphaFold v3. The expected position error graphs from those predictions depict the distance between the same residue across predicted dimer models. The smaller the expected distance between residues indicates greater consistency between different models generated during the prediction.

**(B)** Predicted dimers of Cytb561a consistently align with human DCYTB dimers. The top three predicted models of Cytb561 paralogs from *S. rosetta* (grey structures) were aligned to the crystal structure of human DCYTB. Alignments were restricted to only one chain from each dimeric complex, allowing the second chain to be positioned based on the predicted structure. To better visualize the position of each dimer, the amino-terminal residue is colored in green and the carboxy-terminal residue in magenta. In these alignments, all three models of Cytb561a from *S. rosetta* consistently align with the DYCTB dimer. In contrast, the second polypeptide chain from Cytb561b and Cytb561c display variable positions across all three models.

**Figure S19:** The lumenal surfaces of Cytochrome b561 feature electrostatic properties that are general features of Cytochrome b561 subgroups.

**(A)** The lumenal surface of the DCYTB homodimer (PDB 5ZLG) has negatively and positively charged patches that bind iron by itself or iron complexed with ligands.

**(B)** Loops (yellow) that bridge transmembrane helices form the lumenal surface of DCYTB.

**(C)** Cytb561 subgroups have distinctive electrostatic properties on their lumenal surfaces. The sequences of loops that comprise the lumenal surface of Cytochrome b561 were concatenated to calculate the isoelectric point of the lumenal surface for each paralog. Individual paralogs from each subgroup are displayed as black dots, and the distribution of charges is shown for each subgroup as a grey violin plot. A Kolmogorov-Smirnov test compared the distributions of isoelectric points between Cytochrome b561 subgroups.

**(D)** A character map of lumenal surface charge suggests that electrostatic properties can evolve within Cytochrome b561 subgroups. A maximum likelihood phylogeny of cytochrome B561 proteins (as in Fig. 4A) with the isoelectric points of the lumenal surface for each protein shows that surface properties are variable even in well-resolved clades.

**Figure S20:** Maps of marine upwelling velocities show increased upwelling near coastlines and the equator.

**(A)** Global upwelling velocities at 100m depth averaged over a three year span, measured every 10° latitude and longitude. Missing values are due to shallow locations or landmasses.
**(B)** Upwelling velocities at 100m depth averaged over a three year span at every location sampled during the Tara Oceans Expeditions.

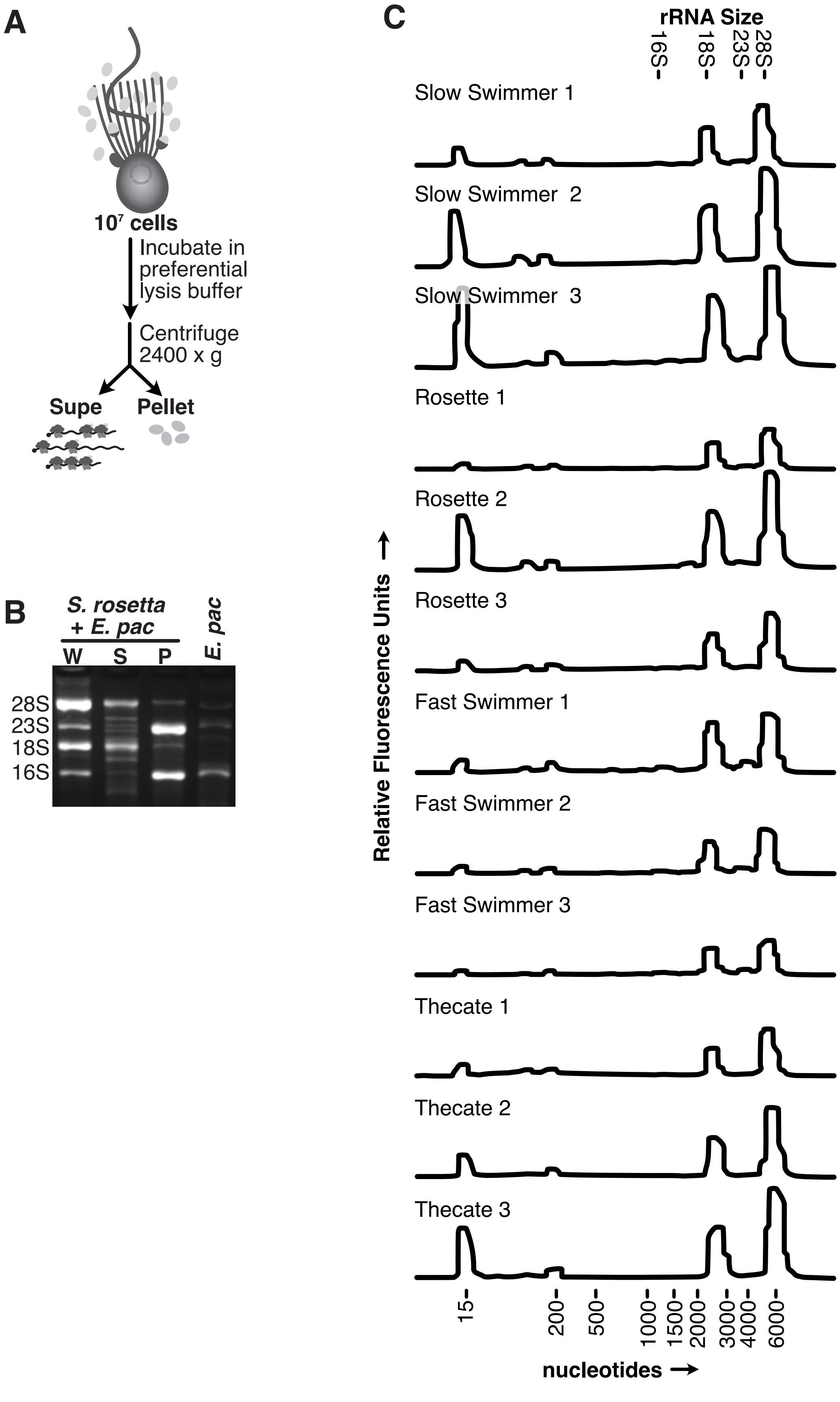


**Figure S1. A preferential lysis buffer enriches RNA from *S. rosetta* to improve transcriptomes.**

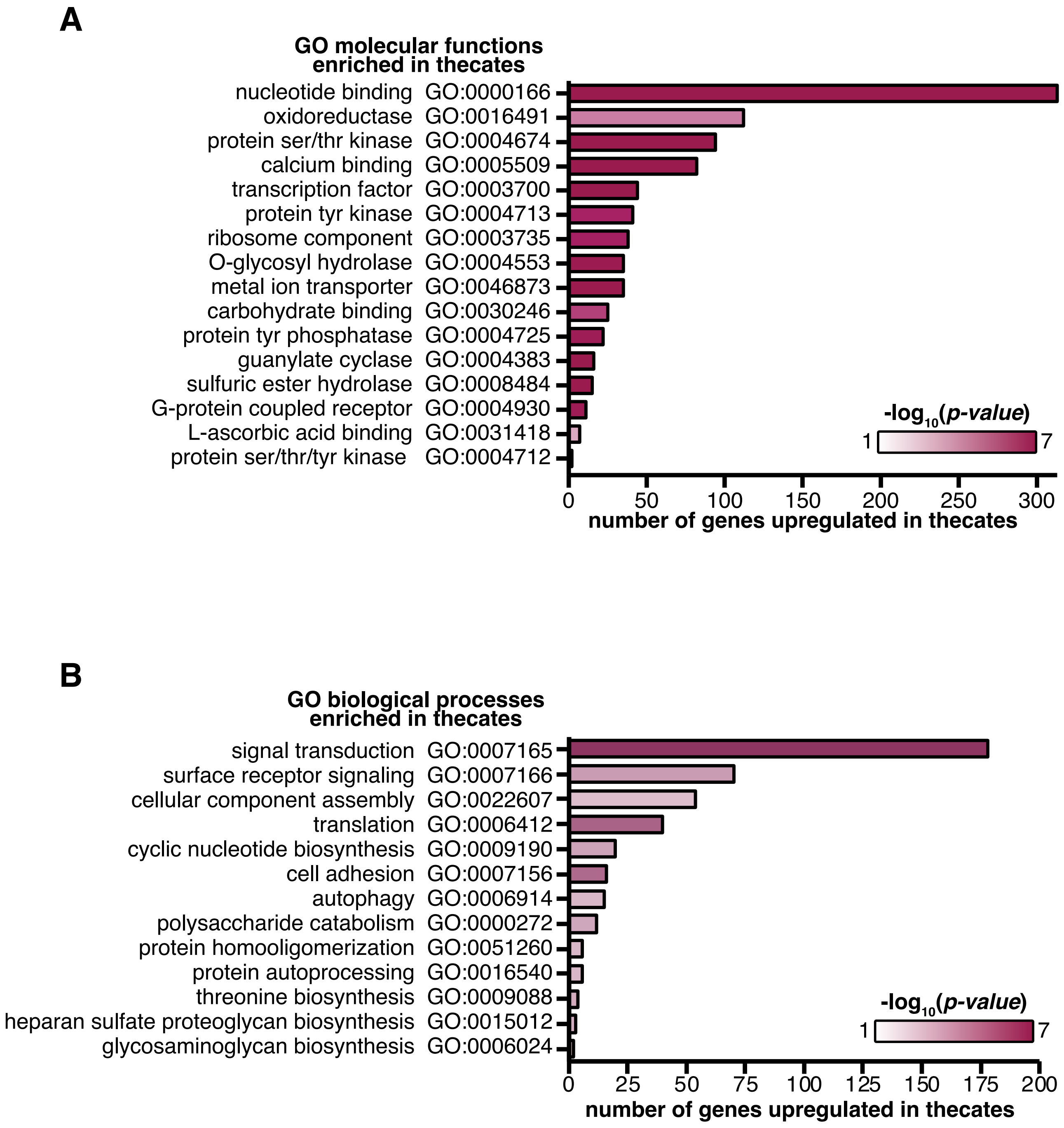


**Figure S2: GO molecular functions and biological processes enriched in thecates.**

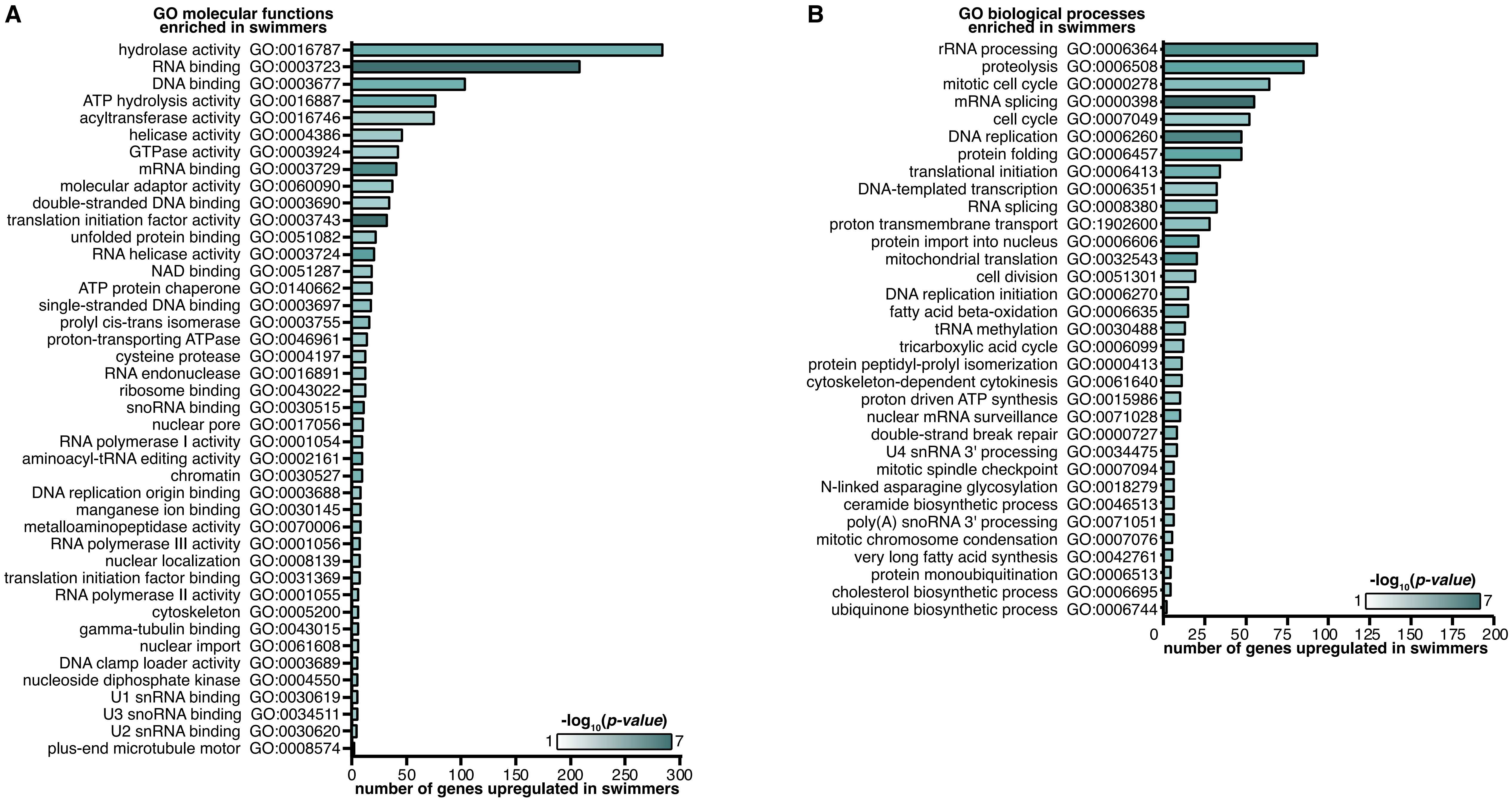


**Figure S3: GO molecular functions and biological processes enriched in swimmers.**

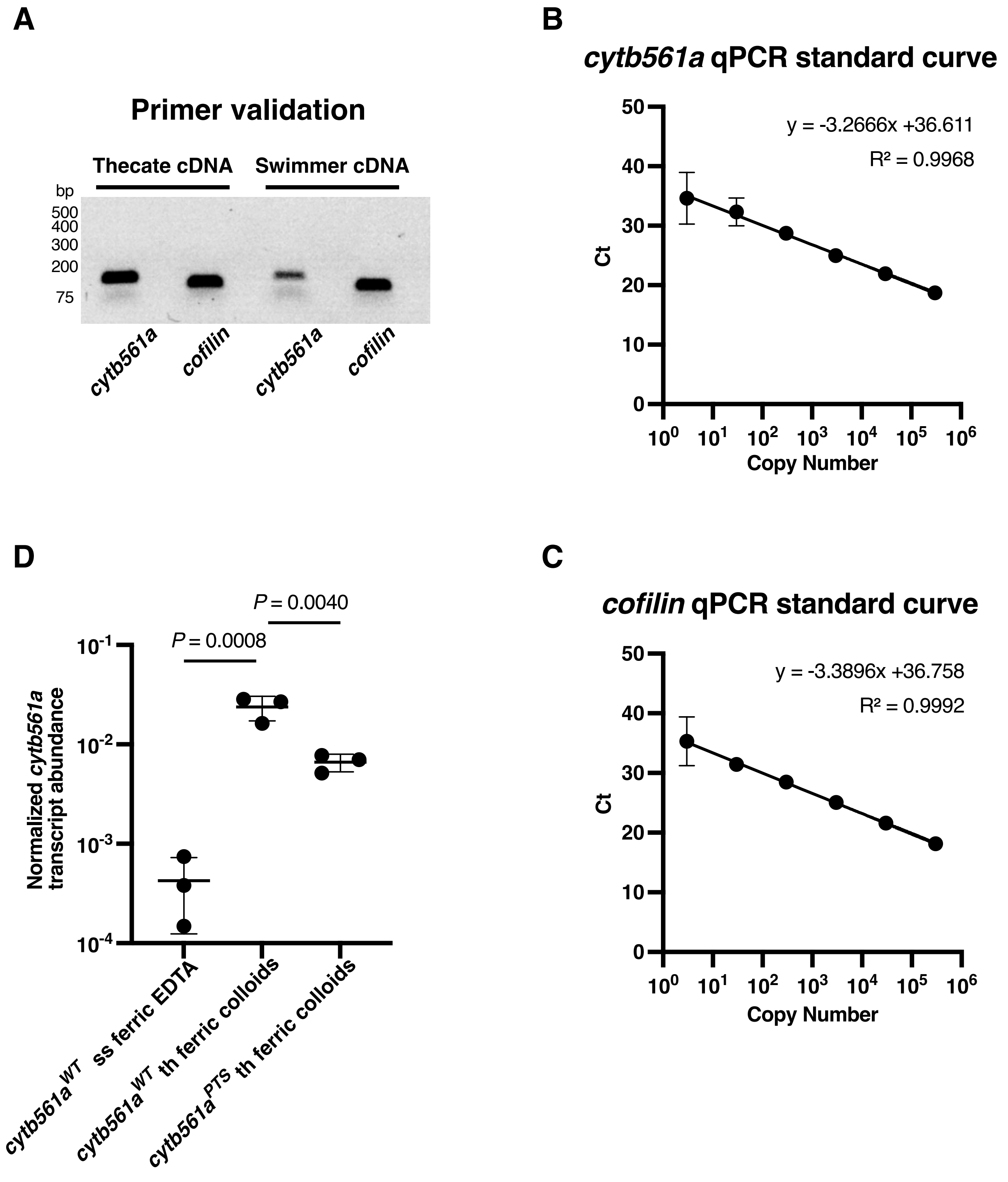


**Figure S4: Validated qPCR primers confirm the disruption of *cytb561a* expression with the *cytb561a*$^{PTS}$ allele.**

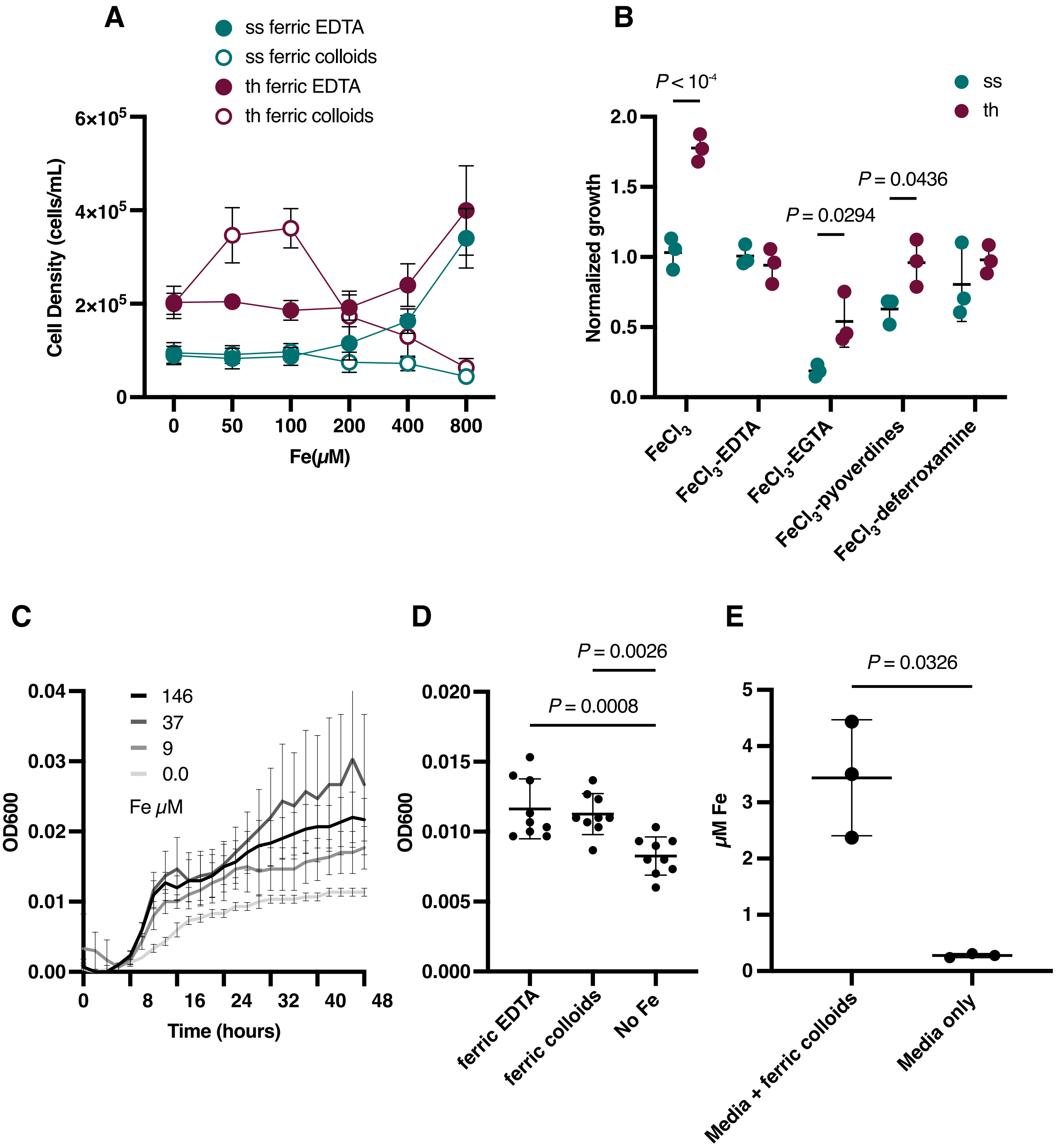


**Figure S5: S. rosetta cell types, but not their feeder bacteria, grow differently with variable iron sources.**

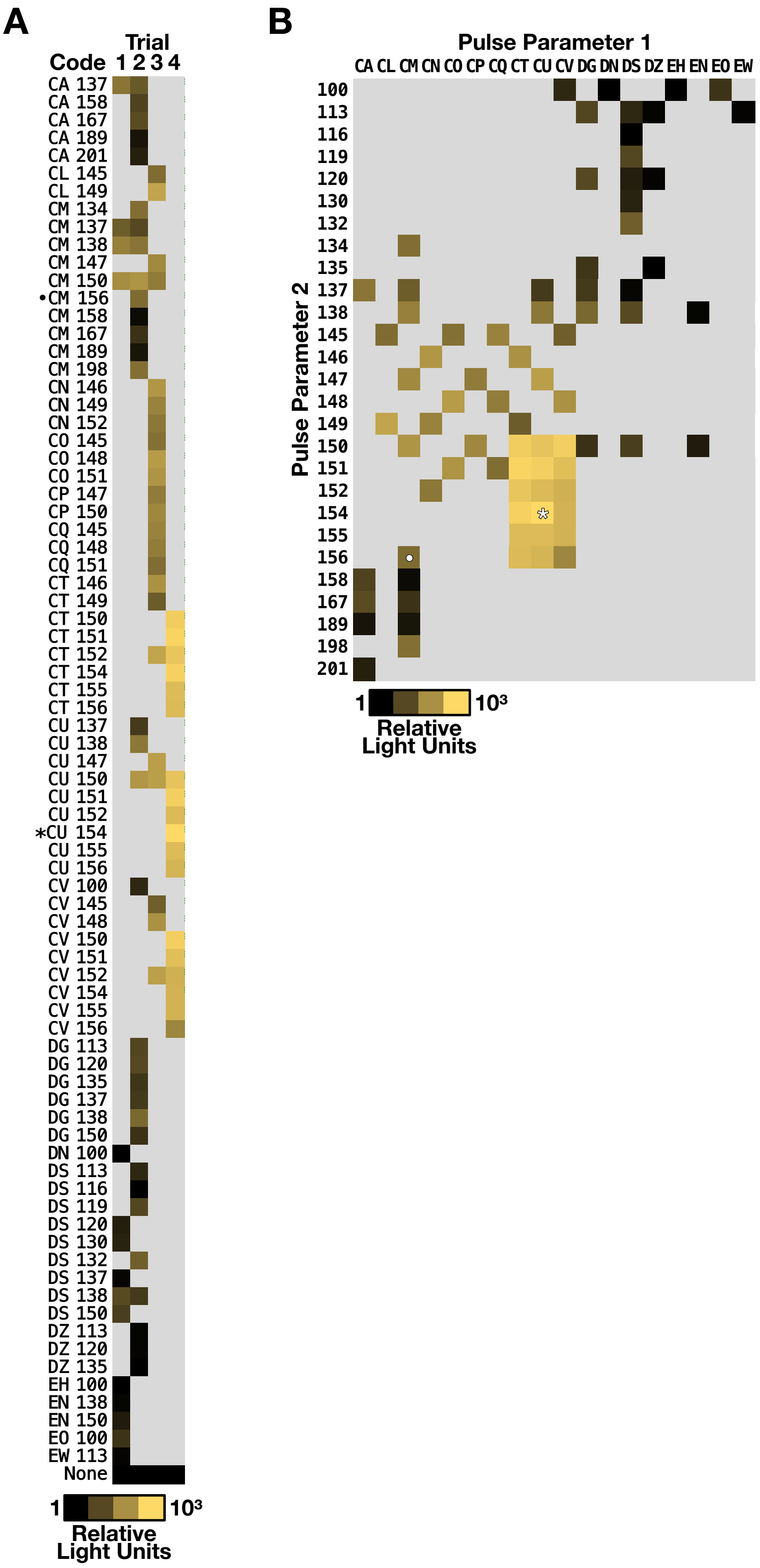


**Figure S6: A screen for optimal nucleofection pulses to improve cargo delivery into *S. rosetta*.**

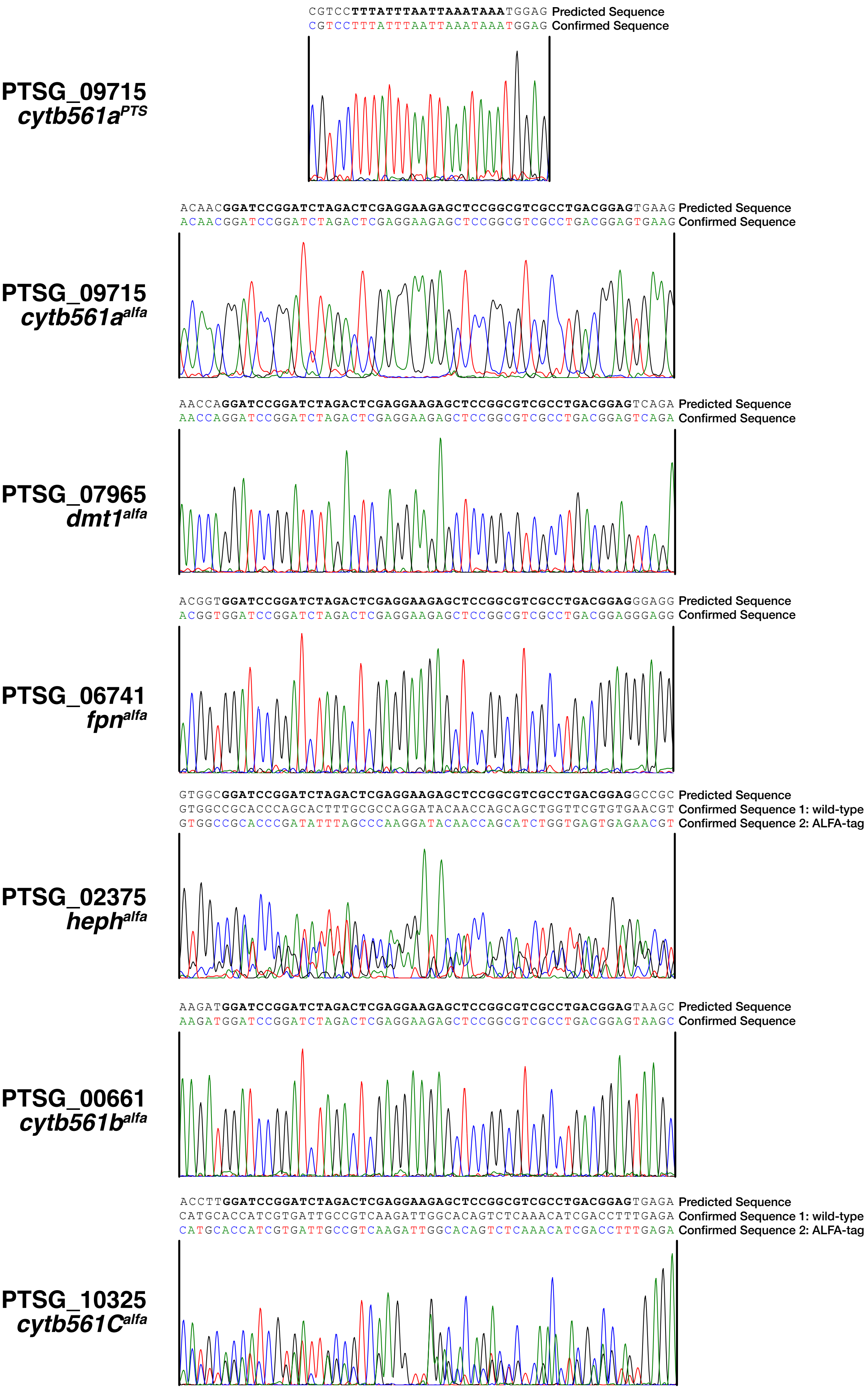


**Figure S7: Genotypes of engineered strains.**

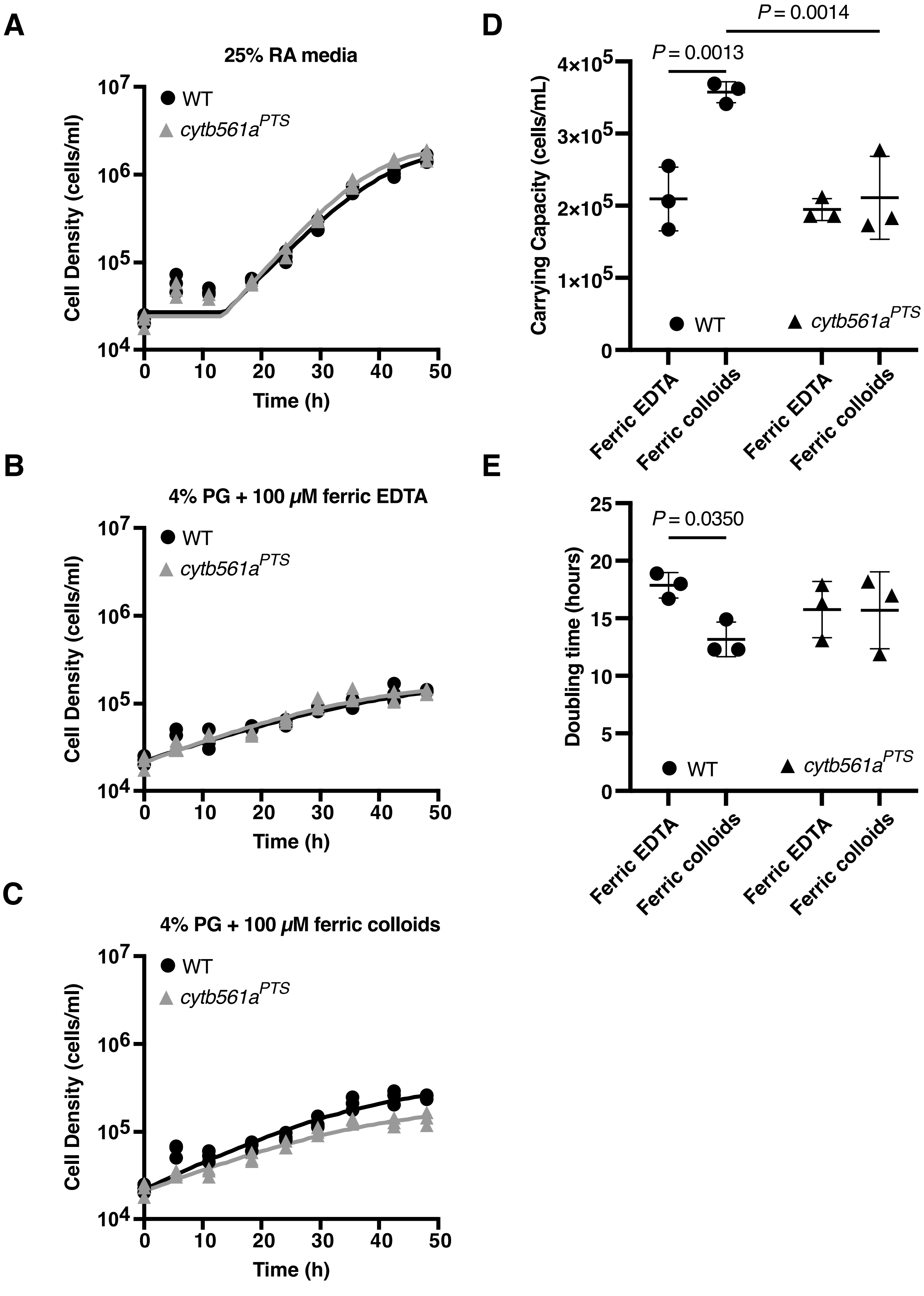


**Figure S8: Thecates grow slight faster and to a larger population size with ferric colloids.**

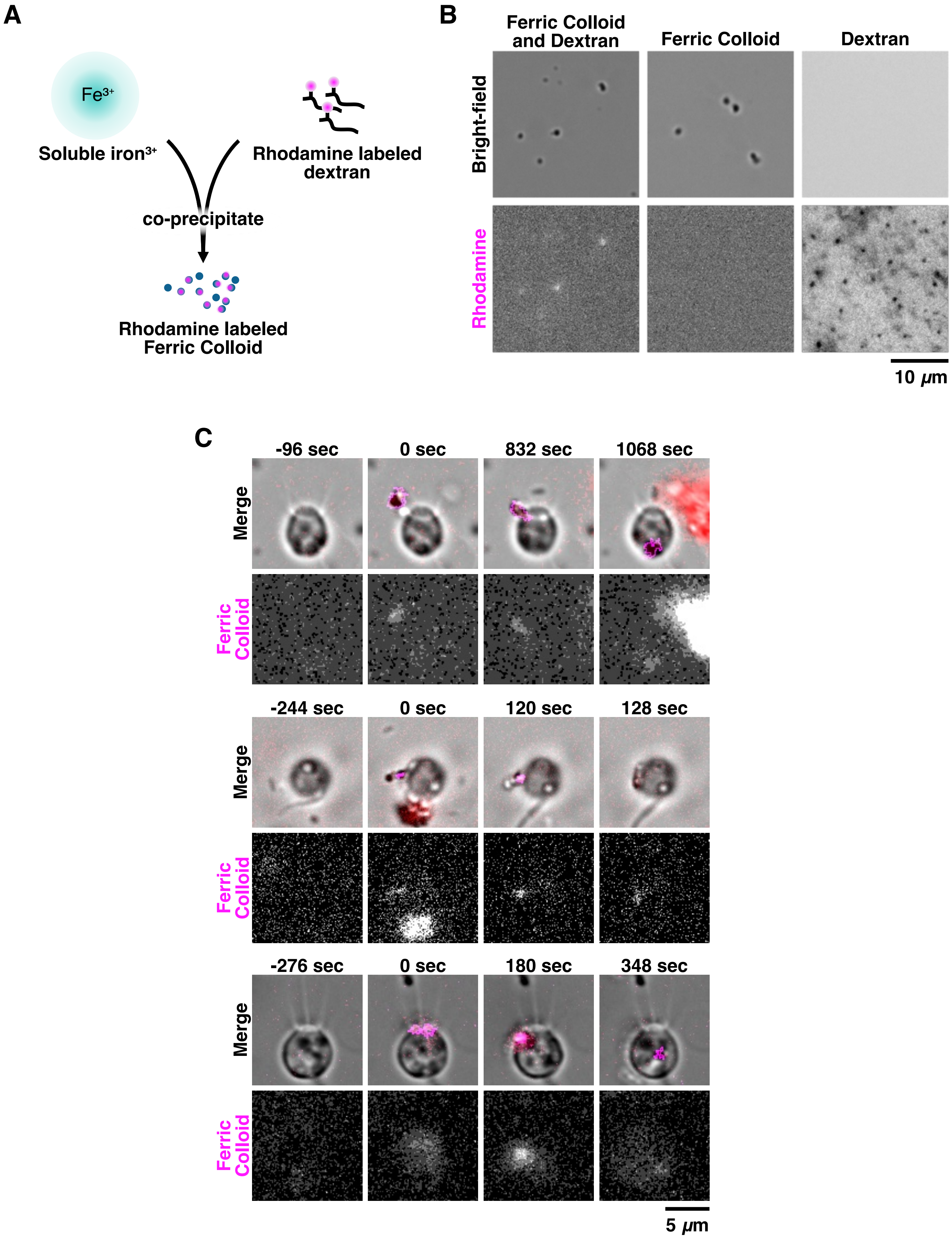


**Figure S9: Thecates phagocytose ferric colloids.**

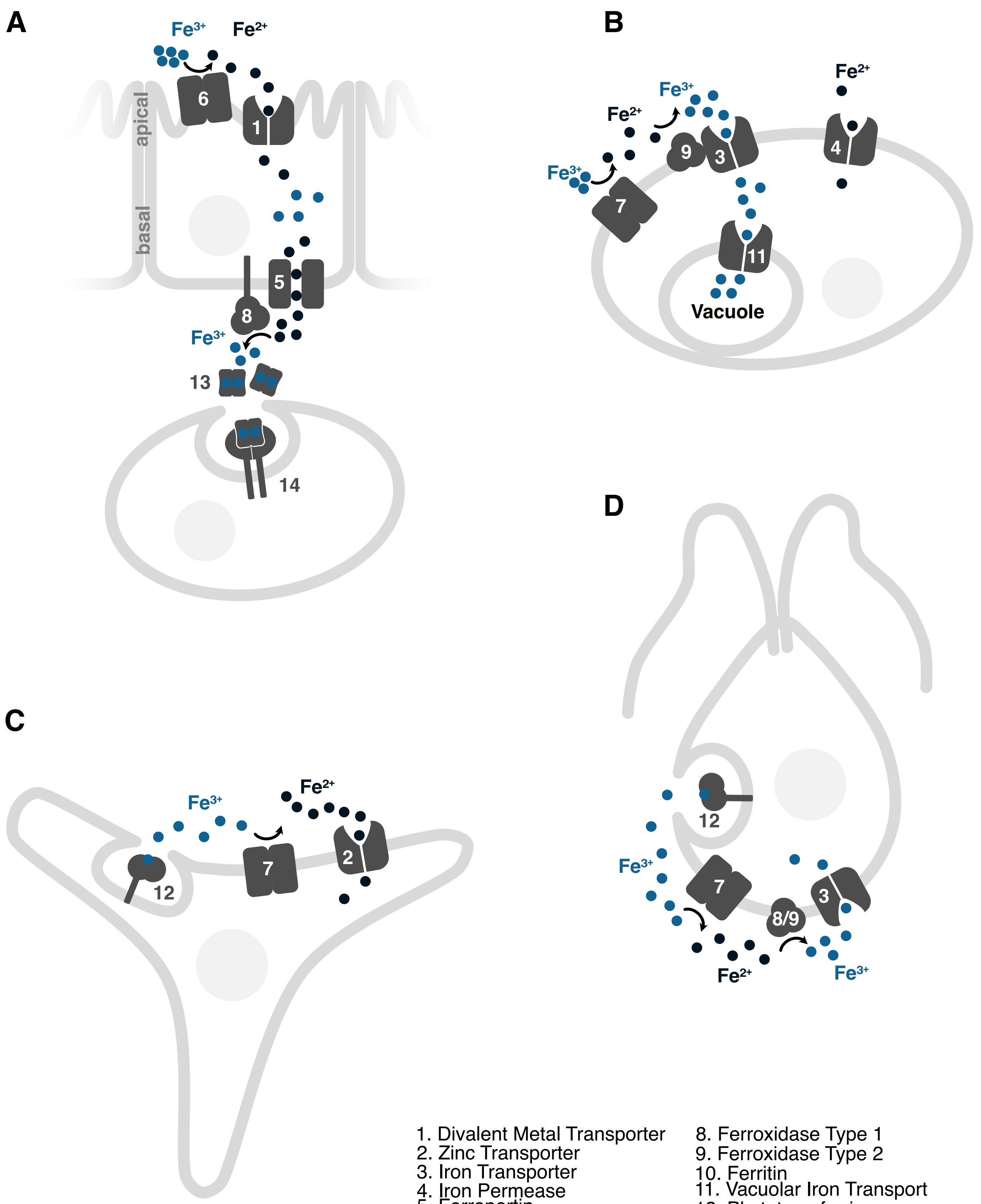


1. Divalent Metal Transporter
2. Zinc Transporter
3. Iron Transporter
4. Iron Permease
5. Ferroportin
6. Cytochrome B561
7. Cytochrome B245
8. Ferroxidase Type 1
9. Ferroxidase Type 2
10. Ferritin
11. Vacuolar Iron Transport
12. Phytotransferrin
13. Transferrin
14. Transferrin Receptor

**Figure S10: Simplified iron acquisition pathways in model eukaryotes.**

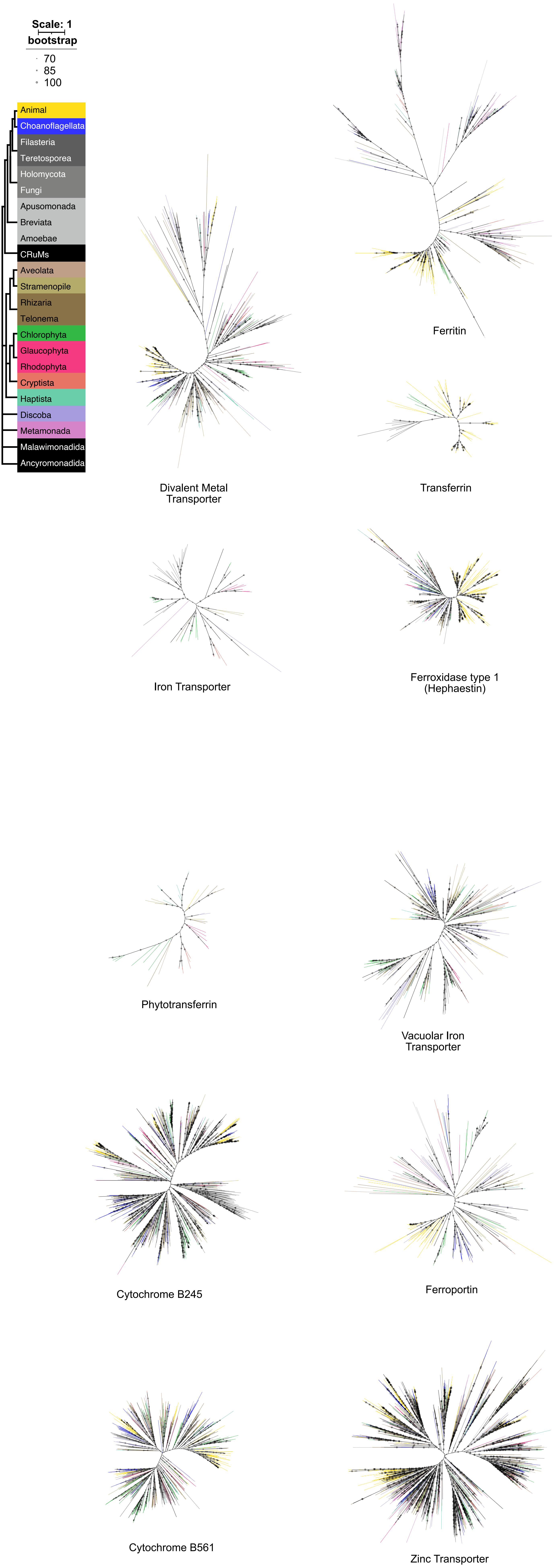


**Figure S11: Phylogenetic trees of iron acquisition proteins.**

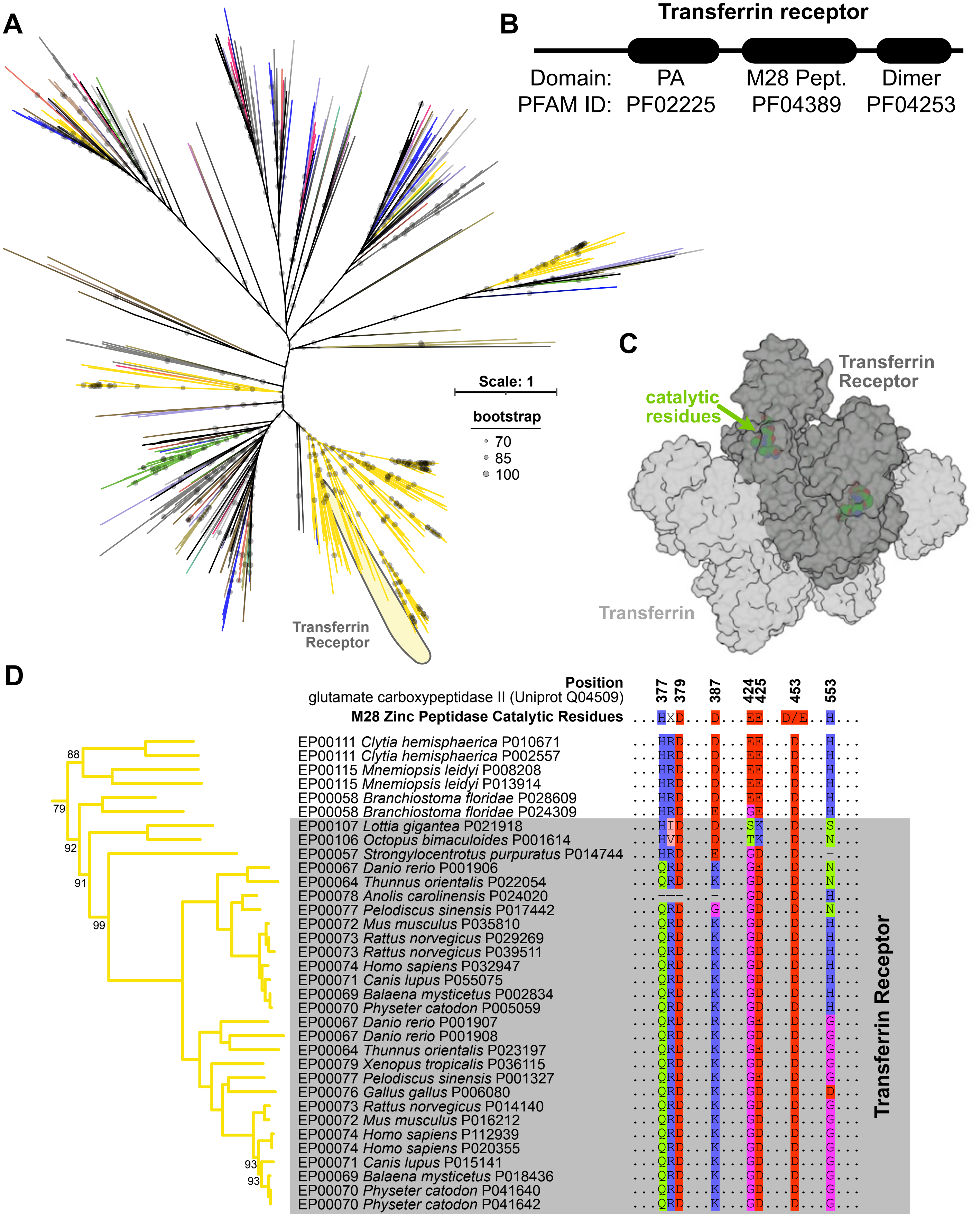


**Figure S12: Animal transferrin receptors evolved from an ancestral M28 peptidase.**

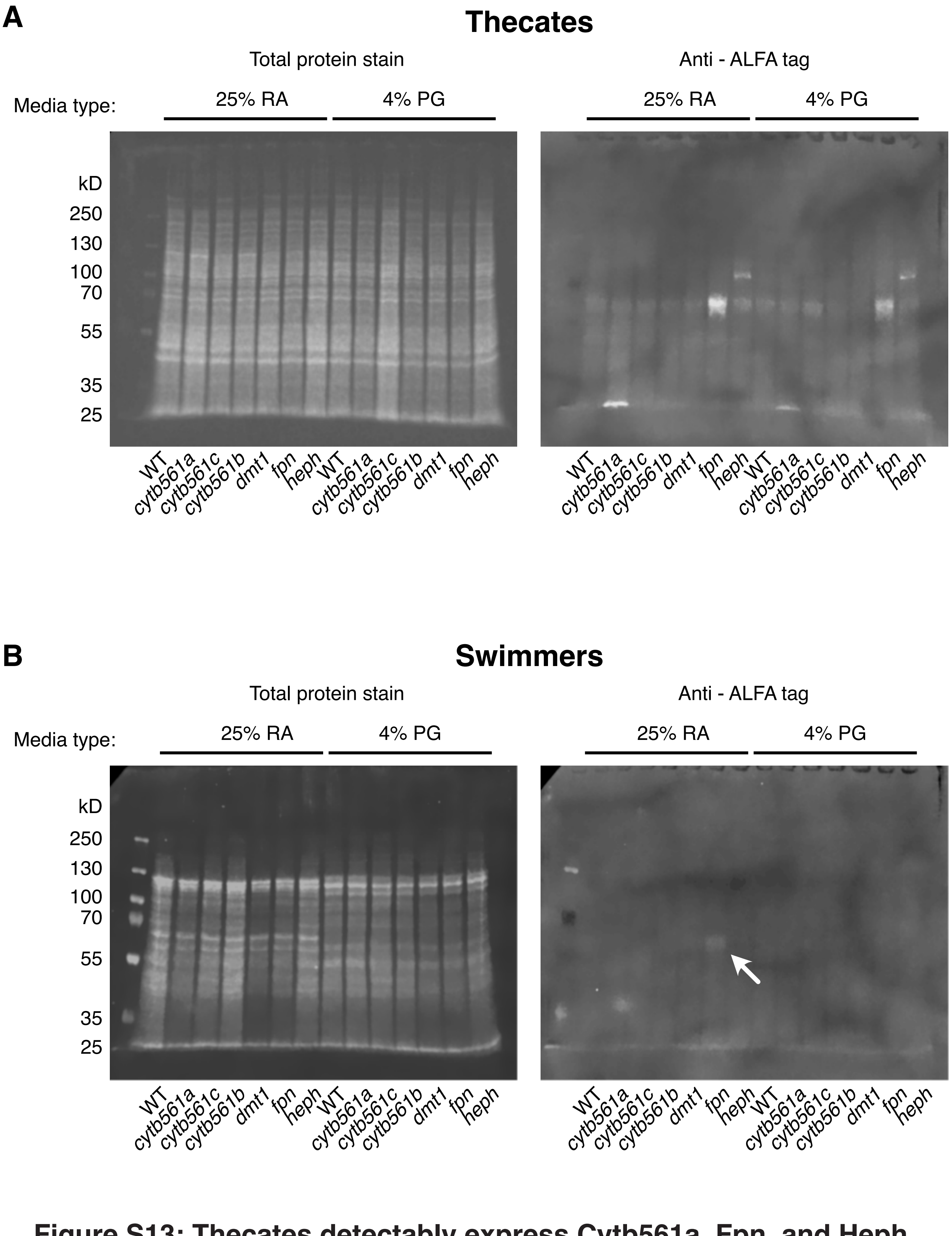


**Figure S13: Thecates detectably express Cytb561a, Fpn, and Heph in western blots.**

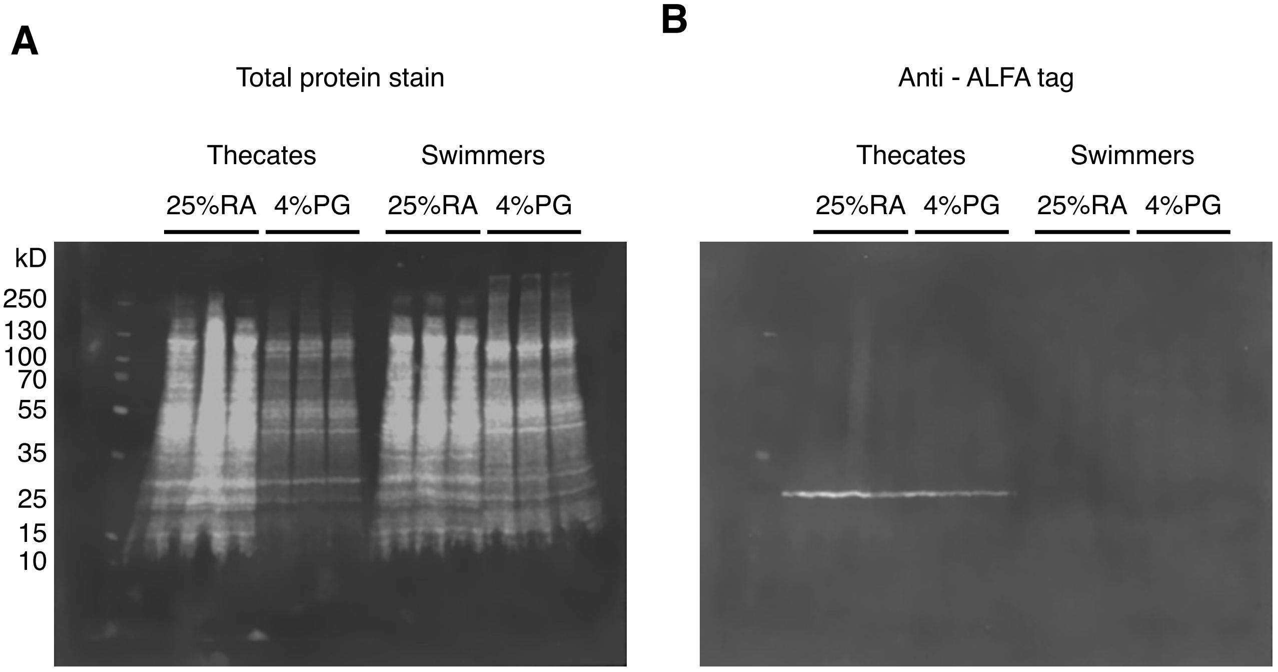


**Figure S14: Thecates stably express Cytb561a independent of iron availability.**

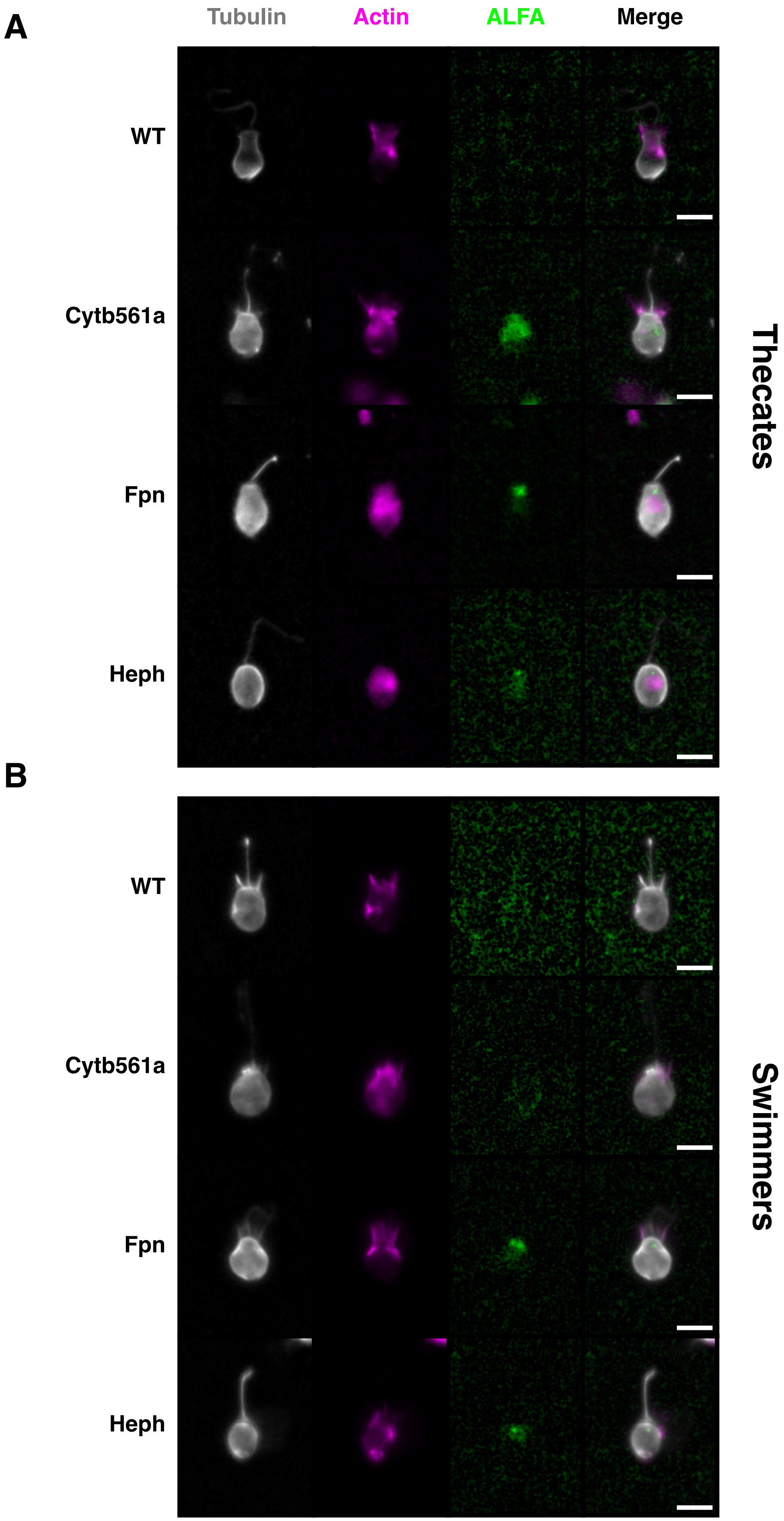


**Figure S15: Additional immunofluorescence of thecates and swimmers.**

A

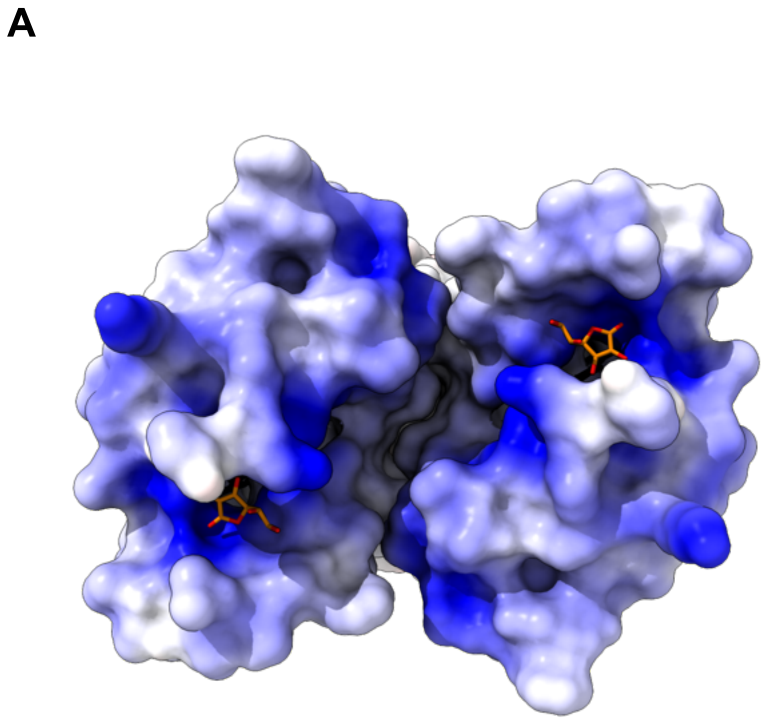

B

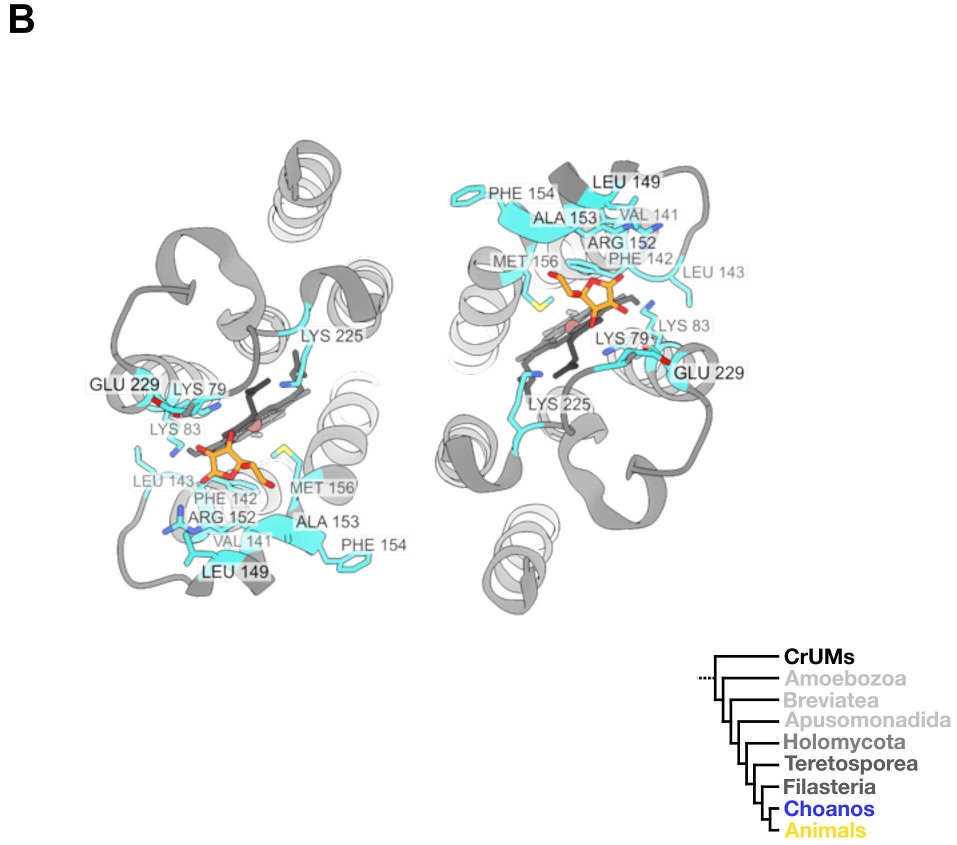


C

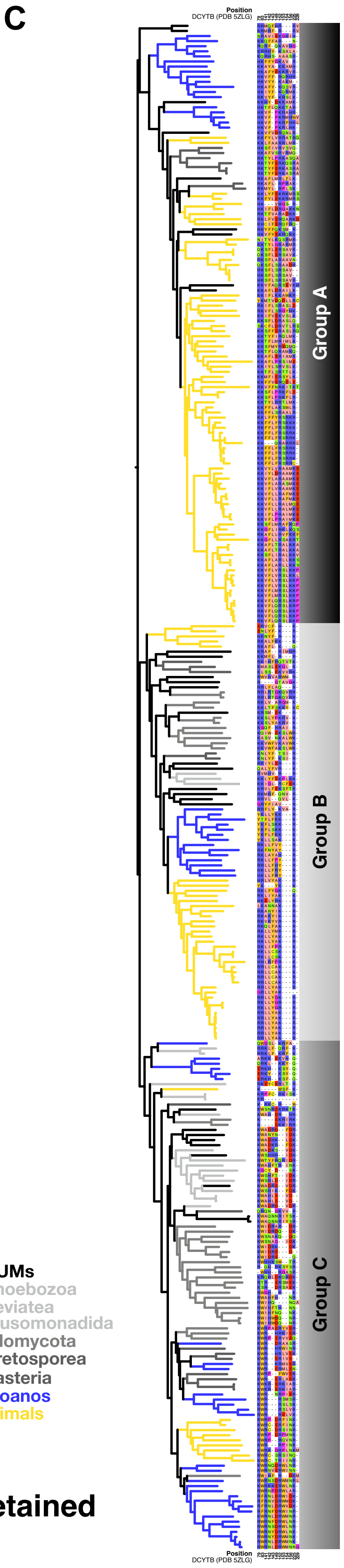


**Figure S16: Cytochrome b561 paralogs have retained ascorbate binding residues.**

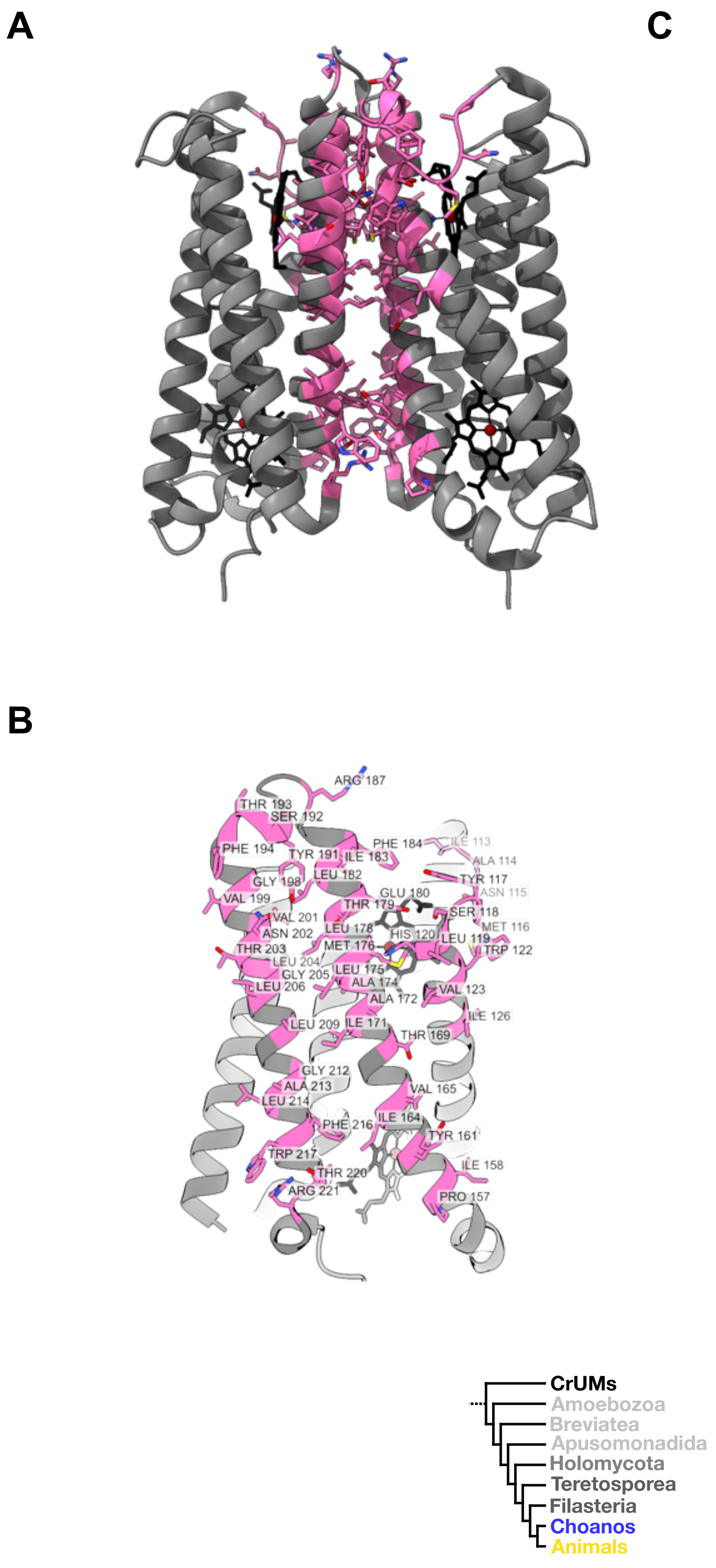

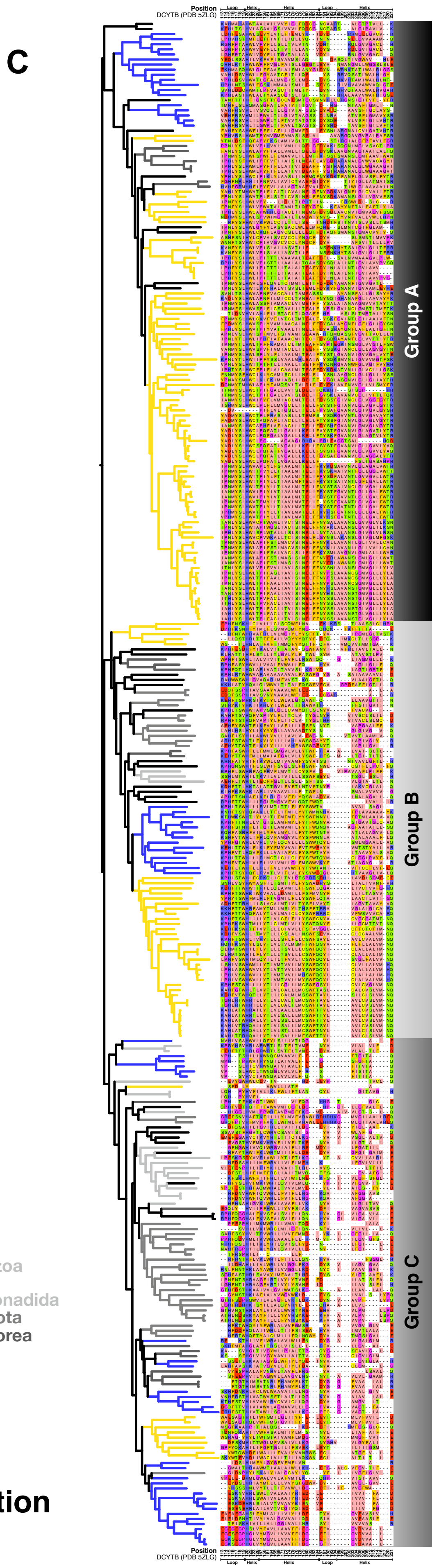


**Figure S17: Cytochrome b561 dimerization is a feature of Group A proteins.**

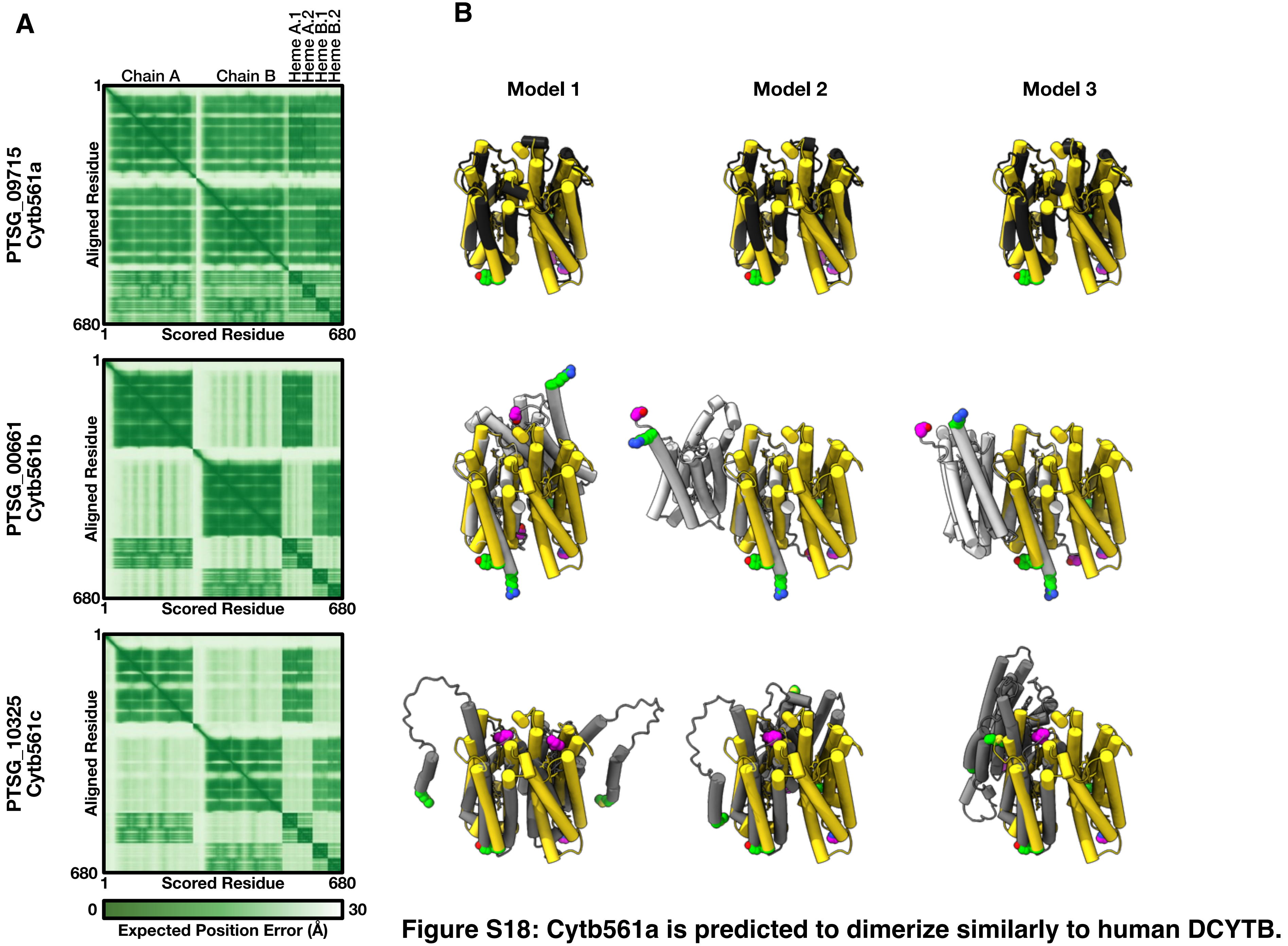


**Figure S18: Cytb561a is predicted to dimerize similarly to human DCYTB.**

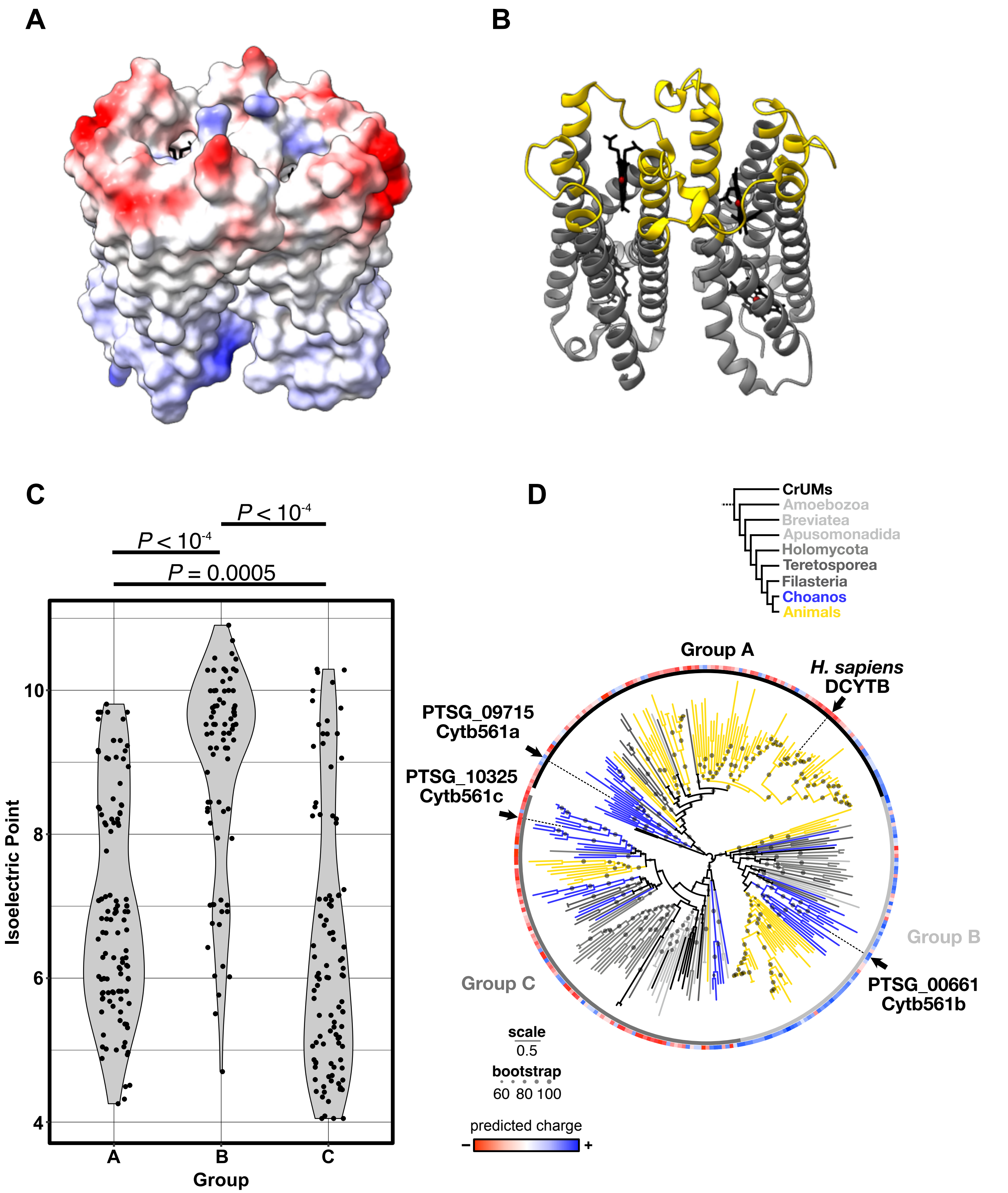


**Figure S19: The lumenal surfaces of Cytochrome b561 feature electrostatic properties that are general features of Cytochrome b561 subgroups**

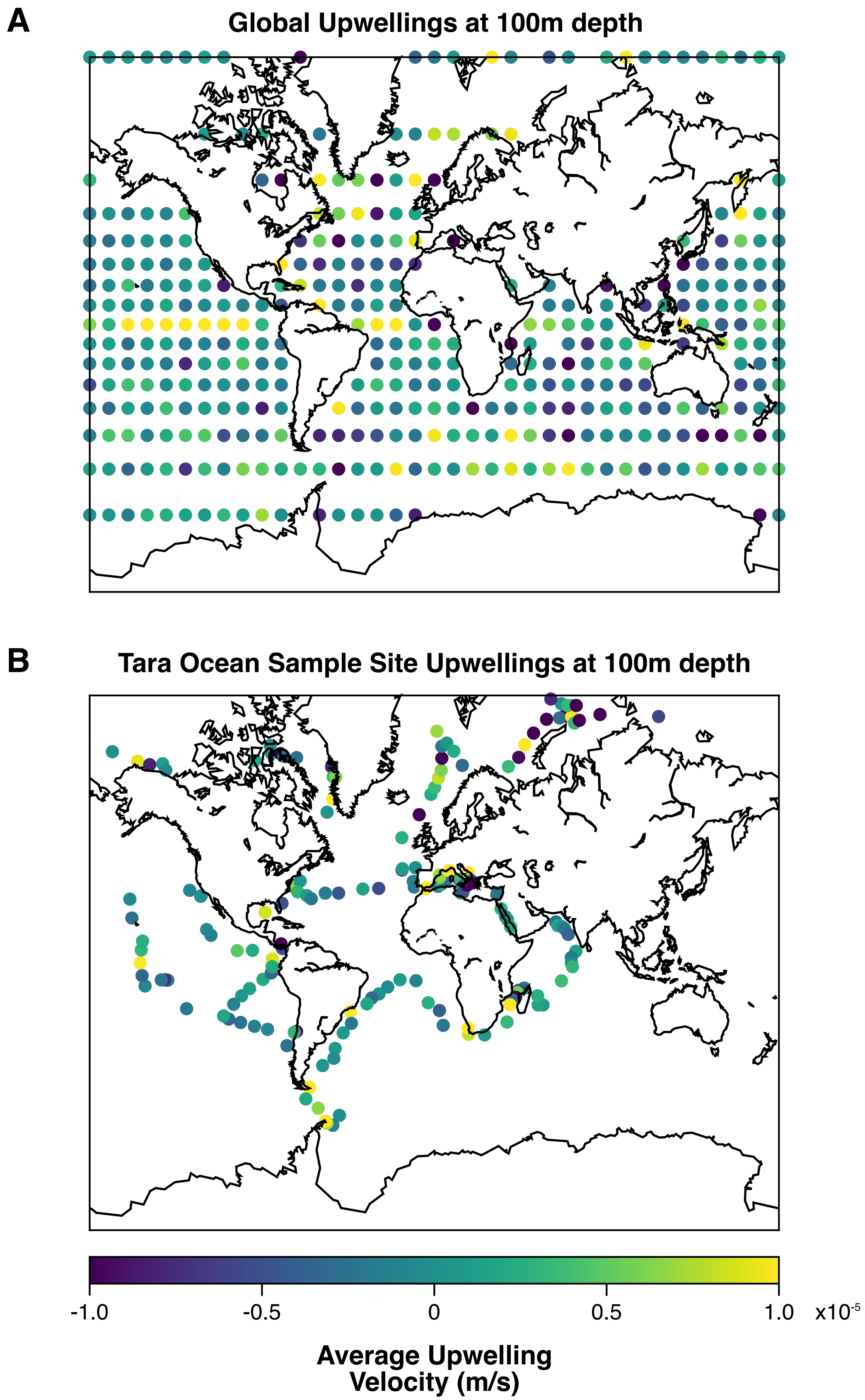


**Figure S20: Maps of marine upwelling velocities show increased upwelling near coastlines and the equator.**

**Tables:**

**Table S1:** Media table.
Components and recipes for all media used: 5% seawater complete (SWC0, 10% cereal grass media (CGM3), 10% red algae (RA), 25% red algae (RA), 15% red algae, 2% peptone, yeast extract, glycerol (15/2), and 4% peptone, glycerol (PG).

**Table S2:** RNA sequencing metadata.
Tabulated strain IDs, culturing conditions, NCBI accession IDs, and sequencing metadata for RNA sequencing.

**Table S3:** Table of differential gene expression between cell types.
Pairwise comparisons of gene expression between cell types, measured by RNA sequencing in transcripts per million (TPM) and scaled reads per base. Each gene's expression is presented as individual triplicate values, along with the mean and standard deviation. Comparisons are made between any two cell types and shown as fold change, standard error, and associated *P*-value and *q*-value. Each gene is also annotated with accession IDs for uniprot, pfam, interpro, orthoDB, unipathway, and gene ontology (GO).

**Table S4:** Table of oligonucleotides and gRNAs for mutant generation and screening.
Tabulated guide RNAs, repair template oligonucleotides, and primers for generating, screening, and isolating mutants.

**Table S5:** Strain table.
The strains of *S. rosetta* used and created during this study, along with the crRNAs, repair templates, Cas12a screening gRNAs, and screening primers for generating each strain.

**Table S6:** Iron acquisition paralog search and phylogenetic tree metadata.
Table of the proteins, HMM models, and PFAM IDs used to search for paralogs, along with the associated protein names used in Fig. 3A. Metadata for HMMer searches, multiple sequence alignments, trimming, and phylogenetic tree are included. The second sheet contains paralog hits filtered for uniqueness and length, which is the raw data used in Fig. 3A. The third sheet is the same paralog heatmap, but without the additional length filtering, resulting in more hits per proteome. Colors of species match Fig. 3A, as does the protein order.
